# Growth-resolved genome-scale metabolic modeling of *Priestia megaterium* SR7 validated by chemostat and ¹³C flux analysis

**DOI:** 10.64898/2026.05.27.728139

**Authors:** Chang Yi Wen Kristy, Yoseb Song, Nathaphon Yu King Hing, Ramanujam Srinivasan Vethathirri, Kristala Prather, Yulan Wang, Janelle R. Thompson

## Abstract

*Priestia megaterium* SR7 is a promising candidate chassis for bioprocess engineering, but its development is limited by the availability of condition-grounded, mechanistic models that can translate experimental measurements into predictive design hypotheses. Here, we present PMSR7, a genome-scale metabolic model for SR7, and evaluate it under a growth-resolved chemostat framework spanning a dilution-rate series. Stable steady states were established across the growth regime, with the highest dilution rate (D = 1.1538 h⁻¹) excluded from growth interpretation due to biomass collapse. Extracellular carbon fluxes were quantified by NMR and reported as mean ± SD, providing an experimental basis for model comparison. PMSR7 was benchmarked using MEMOTE against representative reference reconstructions, supporting structural consistency suitable for constraint-based analyses. Under growth-resolved simulations, ATP demand scaled linearly with growth rate, enabling inference of maintenance-energy behavior across the regime. Growth-dependent feasibility and magnitude of overflow secretion were evaluated for acetate, lactate, and formate using feasible-space analyses, highlighting both agreement and regime-sensitive limitations. Finally, growth-resolved leucine and valine production was assessed in both raw and fold-change space, with experimental means compared against median-based summaries of sampled model distributions to account for feasible-space skew. Together, these results establish PMSR7 as a reproducible, quality-benchmarked platform for SR7 chassis development and provide a framework for iterative experimental integration in non-model organisms, where the dominant challenge is achieving congruence between measured physiology and model-feasible behavior.

## Introduction

Growth rate is a central determinant of microbial physiology, influencing how carbon, energy, and reducing power are distributed between biomass synthesis, maintenance requirements, and by-product formation (Neidhardt et al., 1990; Pirt, 1975; Scott et al., 2010). While growth-dependent metabolic behavior has been extensively studied in canonical model organisms, it remains less defined in alternative production hosts, particularly under steady-state conditions where growth rate can be controlled independently of nutrient depletion (Egli, 1995; Hoskisson & Hobbs, 2005). For organisms being developed as engineering chassis, quantitative understanding of growth-resolved metabolism is necessary to establish a reliable baseline against which metabolic interventions can be assessed (Lewis et al., 2012; Thiele & Palsson, 2010).

*Priestia megaterium* SR7 is a CO₂-tolerant strain originally isolated from formation waters associated with the McElmo Dome supercritical carbon dioxide reservoir (Freedman et al., 2018; Peet et al., 2015). SR7 can germinate and sustain growth under high-pressure CO₂ conditions that are inhibitory to many microorganisms (Peet et al., 2015). Genomic comparison with related strains identified genes associated with stress tolerance, transport systems, and metabolic plasticity that may support growth under CO₂-associated stresses (Freedman et al., 2018).

Physiological characterization under CO₂-rich environments further demonstrated fermentative activity and organic acid production, indicating active central carbon metabolism despite elevated CO₂ exposure (Peet et al., 2015). Collectively, these traits position SR7 as a candidate host for bioprocessing strategies that operate in CO₂-enriched or solvent-extractive environments.

*P. megaterium* SR7 has also been engineered to produce branched-chain alcohols through implementation of a keto-acid-based conversion route in which native branched-chain amino acid biosynthesis supplies keto-acid intermediates that are redirected toward alcohol formation. In this framework, pyruvate enters valine biosynthesis through condensation to 2-acetolactate by acetolactate synthase, followed by reduction and dehydration steps that generate 2-ketoisovalerate. In engineered systems, 2-ketoisovalerate is converted to isobutyraldehyde by a heterologously expressed α-ketoisovalerate decarboxylase (e.g., from *Lactococcus lactis*) and subsequently reduced to isobutanol by an alcohol dehydrogenase. Leucine biosynthesis proceeds from 2-ketoisovalerate through carbon-extension reactions prior to transamination, and the corresponding keto-acid intermediate can likewise serve as substrate for decarboxylation and reduction steps leading to isopentanol formation. Because branched alcohols derive directly from branched-chain amino acid intermediates, quantitative flux through valine and leucine biosynthesis is directly relevant to precursor availability for engineered production (Atsumi et al., 2008; Boock et al., 2019; Connor & Liao, 2008).

Although branched alcohol production has been demonstrated in *P. megaterium* SR7 (Boock et al., 2019), these studies do not resolve how carbon allocation changes across defined growth regimes, nor do they quantify how competition at central metabolic nodes influences precursor availability. The pyruvate node connects glycolysis, the tricarboxylic acid cycle, overflow metabolite formation, and anabolic precursor synthesis, and diversion of pyruvate toward branched-chain amino acid biosynthesis represents a commitment of carbon that can compete with energy generation and by-product secretion. In engineered keto-acid-based alcohol systems, modeling and design studies have highlighted carbon partitioning and redox balance at this node as determinants of pathway performance (Machado & Herrgård, 2014; Zhang et al., 2008). However, a growth-resolved, experimentally anchored analysis of these relationships has not been established for SR7 under steady-state nutrient limitation.

Continuous cultivation provides a framework for isolating the effect of growth rate on metabolic allocation. In a chemostat, the dilution rate fixes the specific growth rate at steady state, allowing substrate uptake, biomass formation, and by-product secretion to be quantified under constant environmental conditions (Monod, 1949; Novick & Szilard, 1950; Pirt, 1975). Unlike batch culture, where nutrient depletion and by-product accumulation confound interpretation, steady-state continuous culture permits direct comparison of physiological parameters across defined growth rates (Egli, 1995; Hoskisson & Hobbs, 2005). These experimentally derived exchange rates provide quantitative boundary conditions compatible with steady-state constraint-based modeling assumptions and enable systematic evaluation of growth-dependent carbon partitioning.

Genome-scale metabolic models (GEMs) provide structured representations of cellular metabolism that can integrate genomic annotation with biochemical knowledge and be interrogated under defined environmental constraints. From early stoichiometric reconstructions, the field matured rapidly after publication of one of the first GEMs for *Haemophilus influenzae* Rd KW20 by Edwards and Palsson (1999), followed by expansion toward increasingly curated and data-informed reconstructions in organisms such as *Escherichia coli* and industrially relevant bacteria (Edwards & Palsson, 2000; Feist, 2007; Gu et al., 2019; Y. K. Oh et al., 2007; Reed et al., 2003). Across applications, GEMs have proven useful for hypothesis generation and metabolic engineering because they provide a consistent accounting framework for mass balance and cofactor usage and allow exploration of feasible pathway usage under explicit constraints (Gu et al., 2019; Orth et al., 2010; Thiele & Palsson, 2010). At the same time, predictive reliability depends on reconstruction integrity and on validation using measurements that directly constrain internal carbon routing rather than only reproducing growth (Lewis et al., 2012; Robaina Estévez & Nikoloski, 2014; Thiele & Palsson, 2010).

Validation is essential in genome-scale reconstruction, particularly for non-model organisms where incomplete annotation, inconsistent metabolite identifiers, and duplicated reaction definitions can introduce hidden degrees of freedom (Lewis et al., 2012; Robaina Estévez & Nikoloski, 2014; Thiele & Palsson, 2010). Standardized identifier systems such as the BiGG Models namespace improve cross-model consistency and reduce ambiguity in reaction and metabolite mapping (King et al., 2016), while benchmarking frameworks such as Memote provide systematic testing of stoichiometric consistency, annotation completeness, and network integrity (Lieven et al., 2020). However, structural compliance alone does not guarantee predictive utility. Demonstrating that a reconstruction produces physiologically consistent behavior under experimentally defined constraints remains a necessary step in establishing its suitability for quantitative analysis.

Here, we develop PMSR7, a stoichiometrically consistent genome-scale reconstruction for wild-type *Priestia megaterium* SR7 by reconciling lineage-appropriate *Bacillus* reconstructions within standardized metabolite and reaction conventions (King et al., 2016; Thiele & Palsson, 2010). The reconstruction was guided primarily by iJA1121, the most recent published genome-scale model of *Priestia megaterium* DSM319 (Aminian-Dehkordi et al., 2019), and by iYO844, the extensively curated reconstruction of *Bacillus subtilis* 168 (Y. K. Oh et al., 2007). Leveraging both a closely related *megaterium* template and a historically validated Bacillus framework enabled retention of lineage-specific metabolic content while maintaining consistency with established reconstruction standards.

We establish a growth-resolved constraint framework using aerobic chemostat cultivation under controlled oxygenated conditions, imposing experimentally measured exchange rates across defined dilution rates as physiological boundary conditions. To further restrict internal solution space beyond exchange anchoring, intracellular flux information derived from ¹³C metabolic flux analysis is incorporated as bounds on central carbon reactions. Model performance is then evaluated by comparing predicted leucine and valine production fluxes to independently measured total amino acid production across growth regimes, providing a growth-resolved assessment of physiological consistency. This experimentally constrained aerobic framework establishes the first genome-scale model for *Priestia megaterium* SR7 and defines a quantitative baseline for downstream predictive application, including extension toward CO₂-relevant cultivation environments.

## Methods

### Chemostat cultivation

*Priestia megaterium* SR7 was cultivated in laboratory-scale chemostats with a 260 mL working volume under aerobic, carbon-limited conditions using modified Furch medium supplemented with 1 g·L⁻¹ glucose as the sole carbon source. Cultures were maintained at 37 °C with continuous agitation and sterile aeration supplied at 1.5 L·min⁻¹ via 0.22 µm filtered air. Samples are collected after four hydraulic retention times are reached and the chemostat is at steady state.

Specific uptake and secretion rates were calculated from concentration differences between influent and effluent streams, multiplied by the dilution rate and normalized to biomass concentration.

### Nuclear magnetic resonance spectroscopy (NMR)

Extracellular glucose and overflow metabolites (acetate, lactate, ethanol, and formate) were quantified using nuclear magnetic resonance spectroscopy. Filtered chemostat supernatant (540 µL) was mixed with 60 µL phosphate buffer (Bruker, USA), vortexed thoroughly, and transferred into 5 mm NMR tubes. Spectra were acquired using a III HD 600 MHz Ascend NMR spectrometer (Bruker, USA), with a 5 mm BBI 600 MHz Z-gradient probe and BOSS III shimming system. A one-dimensional NOESY presaturation pulse sequence was employed for water suppression. PBS was used as the solvent, with a sample jet temperature of 5°C, a 1D NOSEY & JRES acquisition pulse sequence and a 26.85°C probe temperature.

NMR spectra was first normalised using total normalisation method, then uploaded to SIMCA for Multivariate analysis. Principal Component Analysis was performed to evaluate sample clustering behavior. Pair-wise Orthogonal partial least squares discriminant analysis (OPLS-DA) models were applied to compare the detailed metabolic changes of selected groups. Permutation test and CV-ANOVA were tested for each model to evaluate and validate the qualities of the OPLS-DA models.

### Liquid chromatography–mass spectrometry (LC–MS)

Extracellular amino acids and intracellular central carbon metabolites were quantified using LC–MS. Cell pellets were lysed using Qiagen (Germany) PowerBead tubes for 10 min on a vortex mixer. Amino acids were derivatized using the 6-aminoquinolyl-N-hydroxysuccinimidyl carbamate-based method. Cell extracts were derivatized using the Waters (USA) AccQ-Tag Ultra Derivatization Kit following manufacturer instructions, then separated using an Agilent (USA) Zorbax Eclipse XDB-C18 column with water containing 0.1% formic acid as mobile phase A and methanol as mobile phase B (Huang et al., 2019). Detection was performed using an AB SCIEX 6500 QTRAP operating in multiple reaction monitoring mode, targeting the characteristic 171 m/z fragment of the derivatized aminoquinoline moiety.

Tricarboxylic acid cycle intermediates and organic acids were derivatized using a 3-nitrophenylhydrazone and 1-ethyl-3-(3-dimethylaminopropyl) carbodiimide protocol and analyzed for magnetic resonance microscopy (MRM) transitions on a Waters (USA) XEVO TQ-S triple quadrupole mass spectrometer (Han et al., 2013).

Central carbon metabolites, including sugar phosphates, were measured without derivatization on an Acquity UPLC BEH Amide column with a 95% H_2_O and 5% acetonitrile, 20 mM ammonium formate and 20 mM ammonium hydroxide mobile phase A, and a 5% H_2_O and 95% acetonitrile mobile phase B. MRM transitions were detected using an AB SCIEX 6500 QTRAP system (Zheng et al., 2024).

For ¹³C labeling experiments, mass isotopomer distributions were extracted from LC–MS measurements of proteinogenic amino acids and central carbon metabolites. Natural isotope abundance correction was performed using IsoCorrectoR prior to flux estimation (Heinrich et al., 2018).

### ¹³C labeling and metabolic flux analysis

For intracellular flux estimation, chemostat cultures were operated under unlabeled conditions for at least four hydraulic retention times to ensure physiological steady state. The feed was then replaced with a glucose mixture consisting of 40% [1,2-¹³C₂] glucose, 20% [U-¹³C₆] glucose, and 40% unlabeled glucose. Cultures were maintained under labeled conditions for one additional hydraulic retention time prior to sampling.

Flux estimation was performed using Metran v3.10 implemented in MATLAB R2012b based on the Elementary Metabolite Unit framework (Antoniewicz et al., 2007; Wiechert, 2001; Zamboni, 2011). Fluxes were estimated by minimizing the variance-weighted sum of squared residuals between simulated and measured mass isotopomer distributions. Statistical adequacy of the fit was evaluated using χ² testing, and 95% confidence intervals were determined by sum-of-squared-residual profiling (Antoniewicz, 2015; Crown & Antoniewicz, 2013).

The biomass formulation used in the ¹³C-MFA model was a reduced central-carbon representation to preserve identifiability of fluxes within the EMU framework. The complete genome-scale biomass reaction was retained separately within the GEM for subsequent integration.

### Genome-scale metabolic model reconstruction

The genome-scale metabolic model PMSR7 was constructed using COBRApy v0.29.1 in Python v3.11.4 (Ebrahim et al., 2013). The annotated *Priestia megaterium* SR7 genome (GenBank accession CP022674.1) was mapped against reference reconstructions iJA1121 from *P. megaterium* DSM319 (Aminian-Dehkordi et al., 2019) and iYO844 from *Bacillus subtilis* 168 (Y. K. Oh et al., 2007). Reactions were curated to ensure stoichiometric consistency, correct reversibility assignments, and elimination of thermodynamically infeasible cycles. Model quality was evaluated using Memote v0.17.0 (Lieven et al., 2020).

### Integration of experimental constraints and growth-resolved simulations

Condition-specific models were generated for each dilution rate by fixing the biomass reaction flux equal to the imposed dilution rate (v_biomass = D). Biomass bounds were set to (D, D). Parsimonious flux balance analysis was applied to obtain minimal-flux solutions under fixed-growth conditions (Lewis et al., 2010).

Prior to application of experimental constraints, all exchange reactions were reset to (0, 1000) mmol·gCDW⁻¹·h⁻¹. Experimentally supported uptake and secretion fluxes were subsequently reopened. Glucose uptake was constrained according to measured uptake envelopes, oxygen uptake was capped at −25 mmol·gCDW⁻¹·h⁻¹, and glucose was enforced as the sole carbon source.

Intracellular ¹³C-derived 95% confidence intervals were mapped to corresponding reactions in PMSR7 and applied as net flux bounds. Bounds were applied incrementally, and feasibility was evaluated after each constraint intersection. If infeasibility occurred, the most recently applied bound was reverted to preserve compatibility among constraints.

All simulations enforced steady-state mass balance (S·v = 0). Linear programming problems were solved using the GNU Linear Programming Kit (GLPK) through the optlang interface in COBRApy (Ebrahim et al., 2013). Flux balance analysis (FBA) was performed by maximizing the biomass objective function subject to imposed chemostat constraints. Parsimonious flux balance analysis (pFBA) was subsequently implemented by minimizing the total sum of fluxes while maintaining the optimal biomass objective value, thereby identifying flux distributions with minimal overall enzyme usage (Lewis et al., 2010). Flux variability analysis (FVA) was performed by solving two linear programs per reaction to determine the minimum and maximum feasible flux values under fixed dilution rate (μ = D), substrate uptake caps, and exchange guardrails, thereby quantifying the admissible flux envelope independent of objective selection. Flux space sampling was performed using the Artificial Centering Hit-and-Run (ACHR) algorithm with 5,000 samples, thinning interval of 200, and random seed of 1 (Kaufman & Smith, 1998).

## Results

### Genome Scale Metabolic model (GEM)

PMSR7 contains 1,196 genes, 1,424 metabolites, and 1,709 reactions, matching the reaction scale of iJA1121 while increasing curated metabolite content and gene associations relative to template reconstruction. Model composition and coverage are summarized in Table 1. This reaction and metabolite coverage provides the biochemical scope required for growth-resolved simulation of central carbon metabolism and branched-chain amino acid biosynthesis.

**Table 1.** Showing comparison of known *Priestia* (previously *Bacillus*) and *Bacillus* models at the time of model construction in comparison to the model built for PMSR7 (*P. megaterium* SR7)

|  | iYO844 | iMZ1055 | iJA1121 | PMSR7 |
| --- | --- | --- | --- | --- |
| Species | *Bacillus subtilis* | *Priestia megaterium* | *Priestia megaterium* | *Priestia megaterium* |
|  | str168 | WSH-002 | DSM319 | SR7 |
| Genome size | 4 215 606bp | 4 983 975bp | 5 097 447bp | 5 449 951bp |
| Genbank acession no. | AL009126.3 | CP003017.1 | CP001982.1 | CP022674.1 |
| Genes | 844 | 1055 | 1121 | 1196 |
| Metabolites | 990 | 993 | 1349 | 1424 |
| Reactions | 1250 | 1112 | 1709 | 1709 |
| Reference | (Y.-K. Oh et al., 2007) | (Zou et al., 2013) | (Aminian-Dehkordi et al., 2019) | This Study |

Structural integrity was then evaluated using Memote benchmarking against iYO844, iJA1121, and iML1515 (Figure 1). All models achieved 100% stoichiometric consistency. PMSR7 substantially improved elemental balance relative to iJA1121, increasing mass balance from 83.9% to 99.7% and charge balance from 82.6% to 91.4%. Overall consistency increased from 50.69% in iJA1121 to 97.15% in PMSR7, approaching curated reference reconstructions (iYO844: 98.23%; iML1515: 98.4%). Annotation coverage in PMSR7 was 75.23% for metabolites and 77.92% for reactions, with 79.98% SBO annotation and a total Memote score of 84. Together, Table 1 and Figure 1 indicate that PMSR7 combines genome-scale coverage with near-reference structural coherence, providing a stable foundation for applying growth-resolved exchange bounds and isotopic constraints in subsequent analyses.

**Figure 1.**
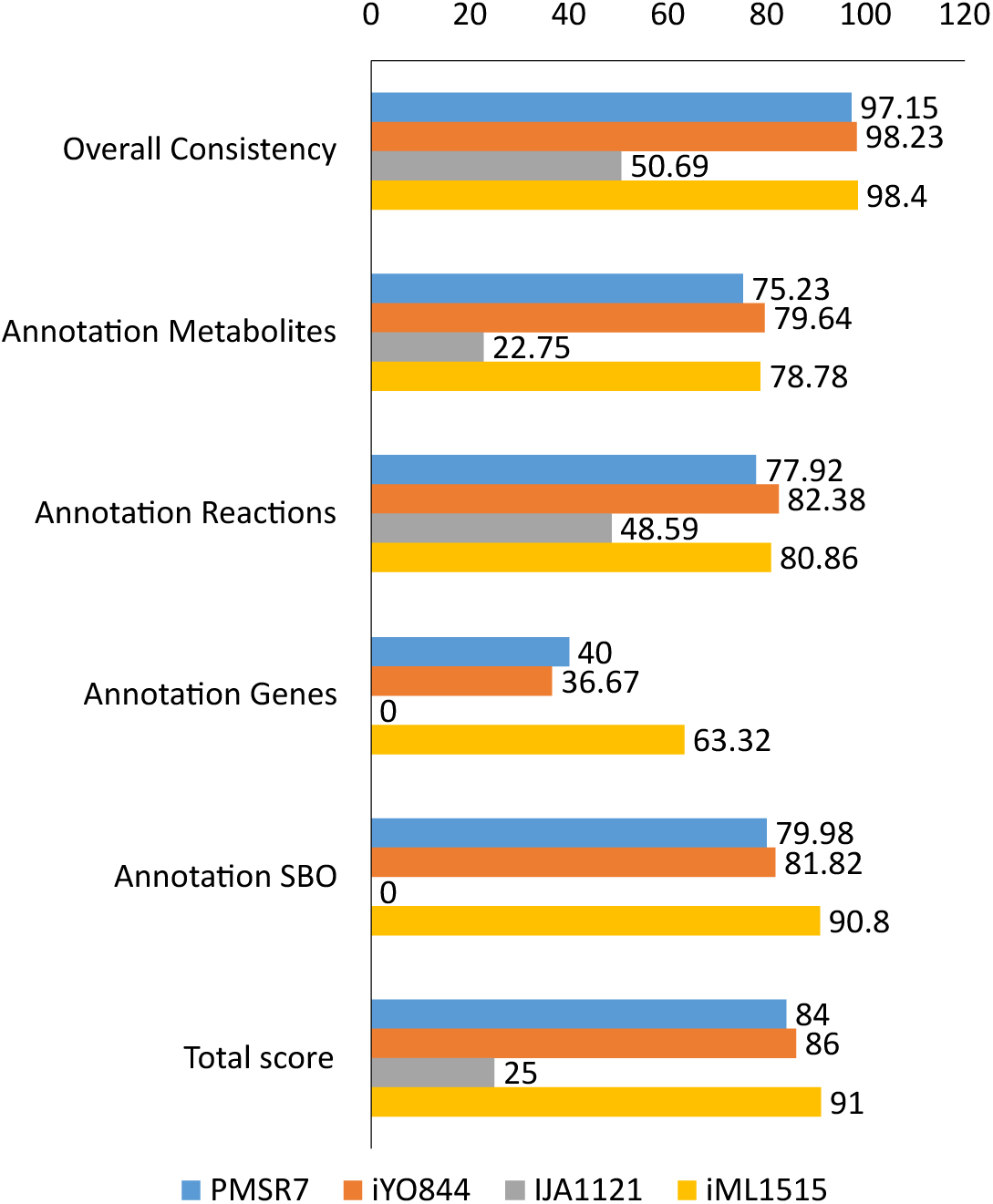
showing the Memote consistency scores of *P. megaterium* SR7 model (PMSR7) against *B. subtilis* str168 (iYO844), *P. megaterium* DSM319 (IJA1121) and *E. coli* str. K-12 substr. MG1655 (iML1515). The overall consistency alongside the Annotation scores contribute to the total score.

### Chemostat-derived growth states define glucose demand and extracellular amino acid phenotypes

Chemostat cultivation generated a defined growth series spanning dilution rates of 0.0231–1.1538 h⁻¹ (Table 2). Because steady-state chemostat growth rate is determined by the dilution rate, this series provided controlled physiological states for downstream model constraint.

Biomass concentration decreased with increasing dilution rate, with the highest flow condition approaching washout-like behaviour (Figure 2A). This highest dilution rate was therefore retained as the upper boundary of the experimental growth series but excluded from subsequent physiological interpretation where biomass collapse would distort biomass-normalized rates.

**Figure 2.**
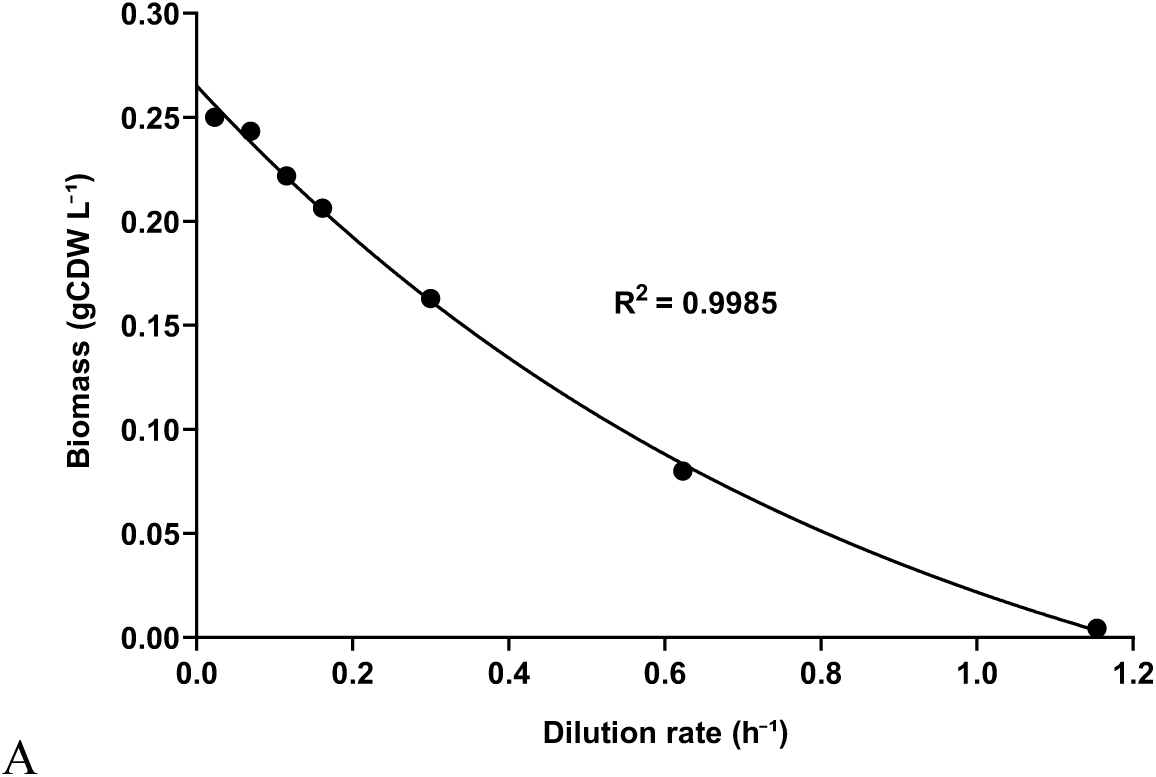
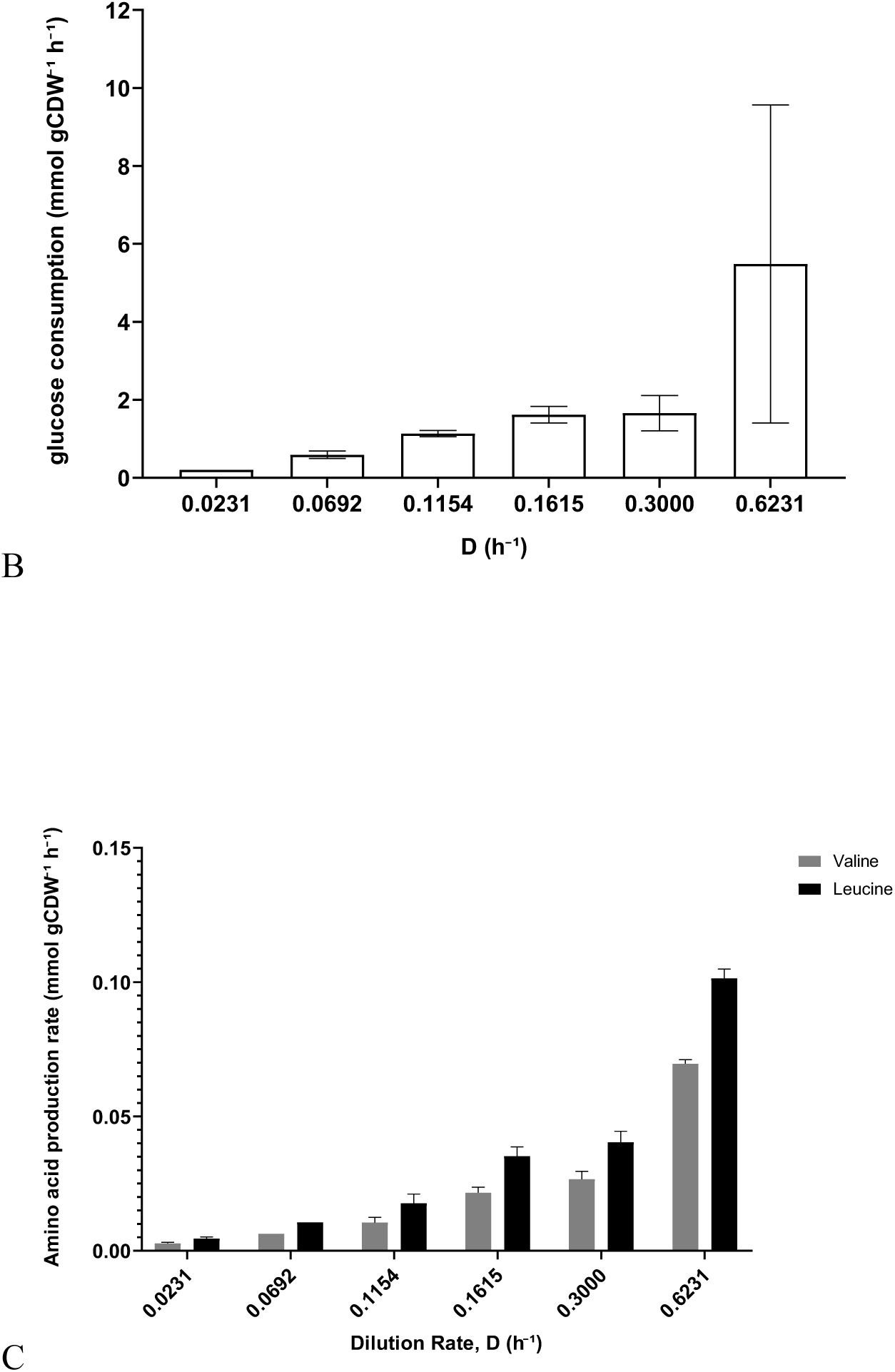
(A) Biomass concentration across dilution rates. (B) Biomass-normalized glucose consumption rate across dilution rates. (C) Biomass-normalized extracellular leucine and valine production rates across dilution rates. Values are shown as mean ± SD where replicate measurements were available. The highest dilution rate was excluded from glucose and amino acid interpretation where biomass collapse prevented reliable normalization.

Biomass-normalized glucose consumption increased with dilution rate, rising from 0.2077 mmol gCDW⁻¹ h⁻¹ at D = 0.0231 h⁻¹ to 5.4851 mmol gCDW⁻¹ h⁻¹ at D = 0.6231 h⁻¹ (Figure 2B). These measured glucose uptake rates provided the primary experimentally derived carbon-input constraints for downstream GEM simulations, ensuring that model predictions were anchored to observed chemostat physiology rather than substrate-unlimited behaviour.

Biomass-normalized extracellular leucine and valine production rates also increased across the interpretable growth range (Figure 2C). These branched-chain amino acids were retained as focused extracellular phenotypes because leucine- and valine-associated metabolism is directly linked to keto-acid precursor availability for branched alcohol production. Unlike glucose uptake, these amino acid measurements were not imposed as intracellular flux constraints but were used as independent physiological readouts for later comparison with sampled model flux distributions.

**Table 1.0.** Flow rates included within the flow rate variation* experiment conducted over 10 weeks Hydraulic Retention Time

| Flow rate, F<br>(ml/min) | Hydraulic Retention Time<br>(HRT)** |  | Dilution rate / Growth rate |  | Doubling time, td |  |
| --- | --- | --- | --- | --- | --- | --- |
|  | (mins) | (H) | (mins<sup>-1</sup>) | (H<sup>-1</sup>) | (mins) | (H) |
| 0.1 | 2600.00 | 43.33 | 0.000385 | 0.0231 | 1802.183 | 30.036 |
| 0.3 | 866.67 | 14.44 | 0.001154 | 0.0692 | 600.728 | 10.012 |
| 0.5 | 520.00 | 8.67 | 0.001923 | 0.1154 | 360.437 | 6.007 |
| 0.7 | 371.43 | 6.19 | 0.002692 | 0.1615 | 257.455 | 4.291 |
| 1.3 | 200.00 | 3.33 | 0.005000 | 0.3000 | 138.629 | 2.310 |
| 2.7 | 96.30 | 1.60 | 0.010385 | 0.6231 | 66.748 | 1.112 |
| 5.0 | 52.00 | 0.87 | 0.019231 | 1.1538 | 36.044 | 0.601 |
| 5.3*** | 49.06 | 0.82 | 0.020385 | 1.2231 | 34.003 | 0.567 |

\*Range of variation determined by the lowest reliable flow rate we can control (0.1ml/min) with high-precision pumps up to the experimentally determined washout rate (5.3ml/min).
\*\*all calculations made with chemostat capacity of 260ml with maximum aeration.
\*\*\*washout flow rate where optical density at 600nm is $0.0 \pm 0.05$

### ¹³C-MFA provides intracellular flux constraints for GEM integration

¹³C-MFA was performed at the 0.7 mL·min⁻¹ chemostat condition to provide intracellular flux information for a representative steady-state growth regime. This condition was selected because it retained stable biomass and measurable glucose consumption while remaining separated from both the slowest-growth and near-washout regimes. The ¹³C-MFA network represented central carbon metabolism, including glycolysis, the pentose phosphate pathway, pyruvate and acetyl-CoA metabolism, and the tricarboxylic acid cycle (Figure 3A). This reduced intracellular network enabled estimation of central carbon fluxes from measured mass isotopomer distributions while preserving the genome-scale biomass formulation separately within PMSR7 for downstream model integration.

**Figure 3.**
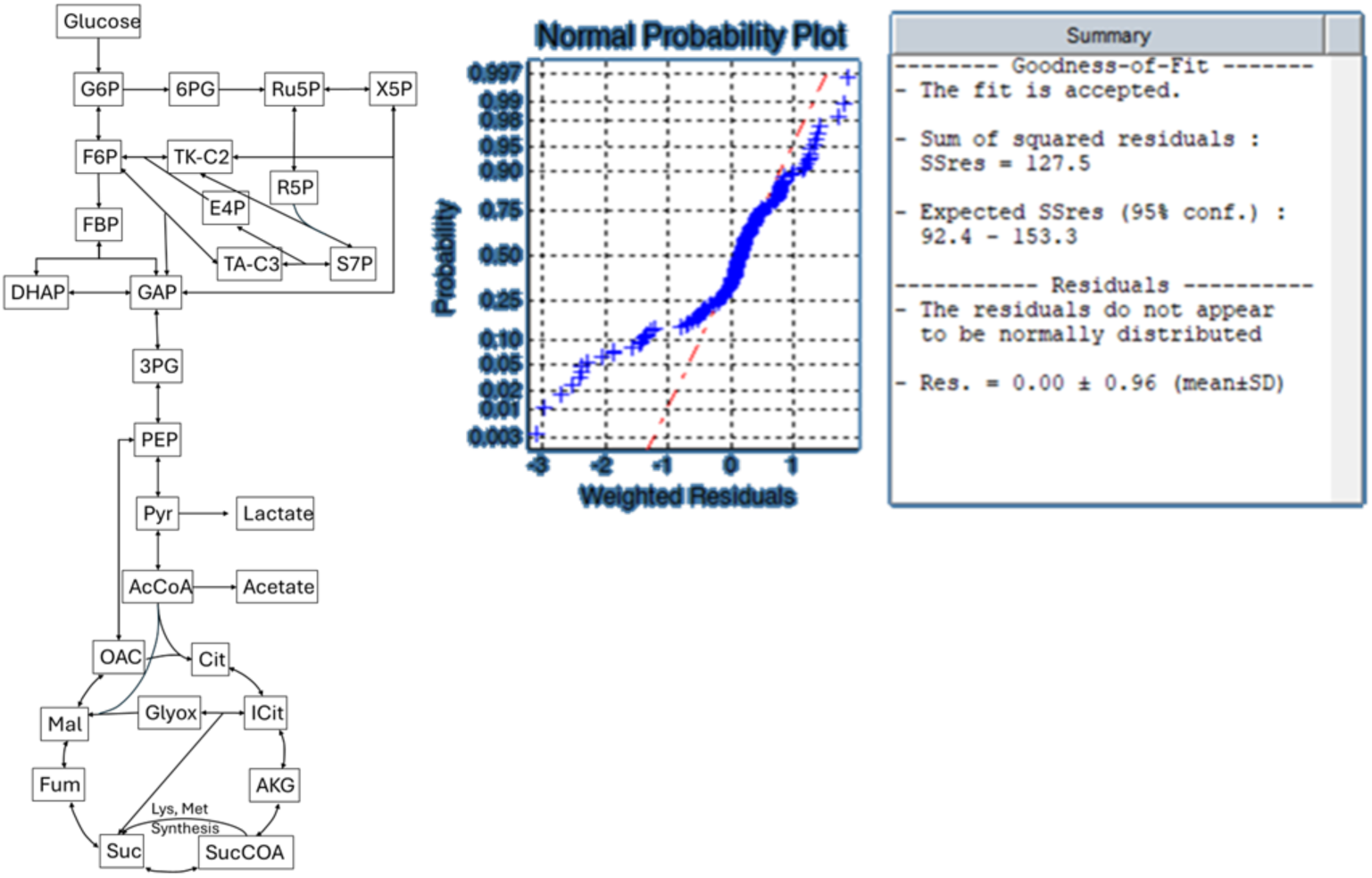
¹³C-MFA network structure and goodness-of-fit assessment used for intracellular flux constraint transfer. (A) Simplified central carbon network used for ¹³C-MFA, showing the represented glycolytic, pentose phosphate pathway, pyruvate/acetyl-CoA, and tricarboxylic acid cycle reactions. This reduced network was used to estimate intracellular central carbon fluxes from measured mass isotopomer distributions. (B) Goodness-of-fit summary and weighted residual probability plot from the accepted ¹³C-MFA solution. The fit was accepted based on the sum of squared residuals falling within the expected 95% confidence interval. The residual probability plot is shown as a diagnostic of weighted residual distribution.

The ¹³C-MFA model achieved an accepted statistical fit, with a sum of squared residuals of 127.5 within the expected 95% confidence interval of 92.4–153.3 (Figure 3B). Although the weighted residuals did not fully conform to a normal distribution, the overall goodness-of-fit criterion supported use of the estimated flux confidence intervals as condition-specific intracellular constraints. These intracellular flux estimates provided information on central carbon routing that could not be resolved from extracellular glucose uptake and amino acid measurements alone.

To integrate this flux information into PMSR7, ¹³C-MFA reactions were mapped to corresponding GEM reactions by reaction equation similarity and applied as net flux bounds using the estimated 95% confidence intervals. Constraint transfer was performed incrementally to preserve feasibility of the genome-scale model. Bounds that caused infeasibility or excessive distortion of the feasible solution space were excluded, while compatible bounds were retained for downstream constrained simulations. The accepted and rejected mapped constraints are provided in Supplementary S1.3. This screening step ensured that the transferred ¹³C-derived constraints remained both experimentally grounded and compatible with the mass-balanced GEM framework.

### Integrated constraint framework supports growth-resolved leucine and valine pathway interpretation

The experimentally constrained PMSR7 framework was next assembled by combining the growth-resolved chemostat measurements, finite aerobic boundary condition, maintenance-energy formulation, and compatible ¹³C-MFA-derived intracellular bounds (Figure 4). This step generated condition-specific models across the dilution-rate series, in which each simulation was anchored to the measured physiological state while retaining genome-scale mass balance.

**Figure 4.**
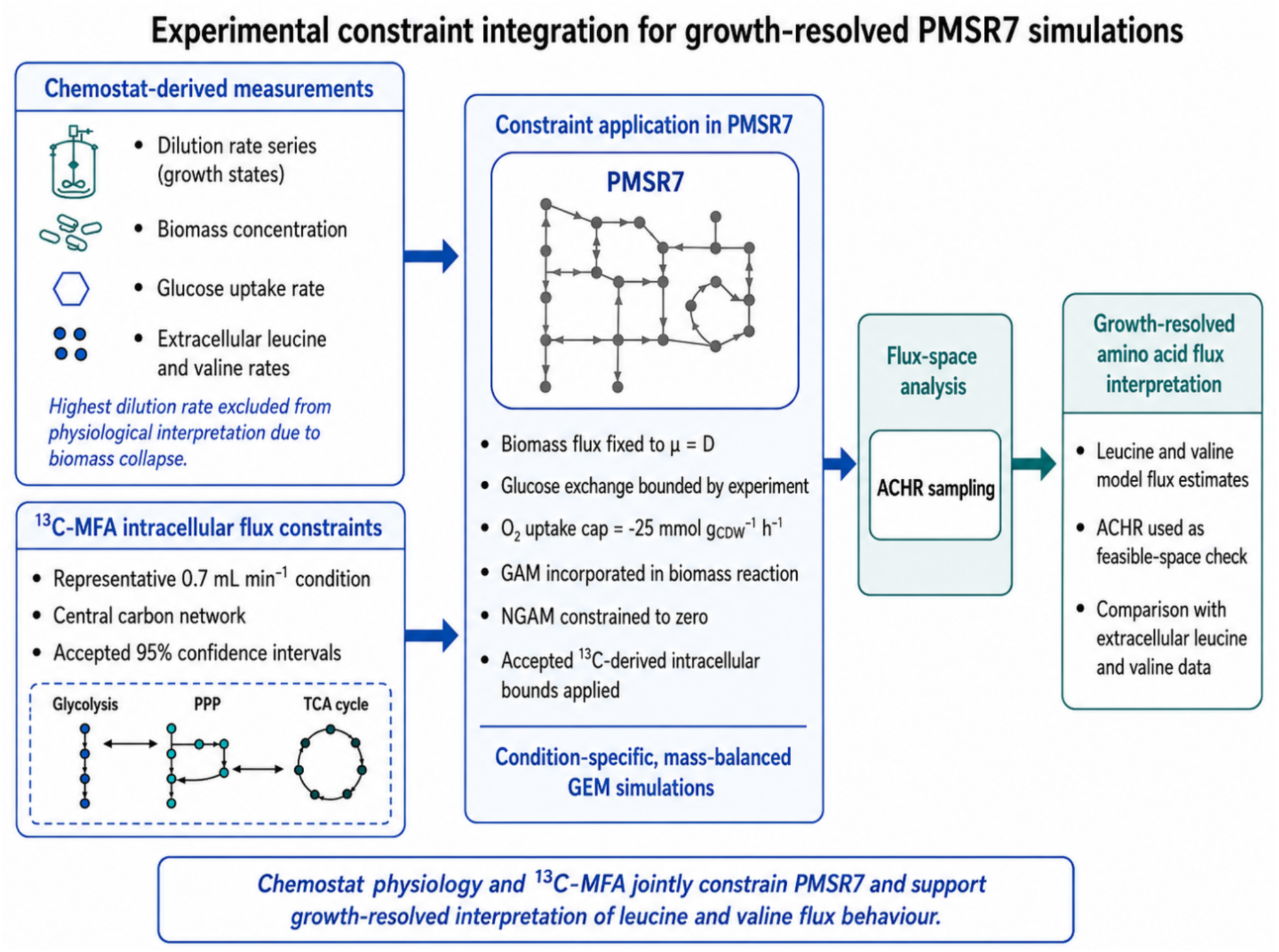
Integration of experimental constraints into growth-resolved PMSR7 simulations. Chemostat-derived measurements provided dilution-rate-defined growth states, biomass-normalized glucose uptake constraints, and extracellular leucine and valine phenotypes. ¹³C-MFA at the representative 0.7 mL min⁻¹ condition provided intracellular central carbon flux bounds. These datasets were integrated into PMSR7 by fixing biomass flux to μ = D, constraining glucose exchange using measured uptake rates, applying a finite oxygen uptake cap, incorporating growth-associated maintenance into the biomass reaction, constraining non-growth-associated maintenance to zero, and applying accepted ¹³C-derived intracellular bounds. The resulting condition-specific, mass-balanced models were analyzed using ACHR sampling as a feasible-space check to support growth-resolved interpretation of leucine- and valine-associated flux behaviour.

For each dilution rate, biomass flux was fixed to the corresponding chemostat growth rate, reflecting the steady-state relationship μ = D. Glucose uptake was constrained using the measured biomass-normalized uptake rates, while glucose was retained as the sole organic carbon source. Oxygen uptake was capped at −25 mmol gCDW⁻¹ h⁻¹ to maintain an aerobic but bounded solution space. As an order-of-magnitude anchor, recent curated *B. subtilis* GEM work supplied oxygen up to approximately 18 mmol gCDW⁻¹ h⁻¹ to match experimental physiology, supporting the use of a finite oxygen uptake boundary rather than an unconstrained aerobic exchange (Dauner & Sauer, 2001; Neal et al., 2024). Growth-associated ATP demand was incorporated through the biomass reaction, and non-growth-associated maintenance was constrained to zero. Accepted ¹³C-MFA-derived bounds were then applied to compatible intracellular reactions following feasibility screening.

ATP demand was incorporated using the carbon-limited growth regime, rather than the full dilution-rate series, because the highest growth conditions were affected by biomass collapse and were not used for physiological interpretation. Under the finite oxygen uptake window of [−25, 0] mmol gCDW⁻¹ h⁻¹, the carbon-limited subset showed a strong linear relationship between modelled ATP maintenance requirement and growth rate (y = 61.987x − 0.3938; R² = 0.9959; Supplementary Figure S2.1B). The slope was therefore applied as the growth-associated maintenance coefficient in the GEM biomass reaction, where ATP hydrolysis terms represent growth-associated ATP cost (GAM = 61.987 mmol ATP gCDW⁻¹). Because the fitted intercept was slightly negative, non-growth-associated maintenance was not imposed as a positive ATP drain; instead, the lower bound of the ATP maintenance reaction (ATPM) was set to zero.

The computational leucine and valine flux estimates increased monotonically with dilution rate across the interpretable growth range (Figure 5). For leucine, the computational estimate increased from 0.0078 mmol gCDW⁻¹ h⁻¹ at D = 0.0231 h⁻¹ to 0.2096 mmol gCDW⁻¹ h⁻¹ at D = 0.6231 h⁻¹, while the corresponding experimental leucine production rate increased from 0.0045 ± 0.0006 to 0.1014 ± 0.0035 mmol gCDW⁻¹ h⁻¹. For valine, the computational estimate increased from 0.0069 to 0.1857 mmol gCDW⁻¹ h⁻¹, while the experimental valine production rate increased from 0.0028 ± 0.0003 to 0.0696 ± 0.0015 mmol gCDW⁻¹ h⁻¹ over the same dilution-rate range.

**Figure 5.**
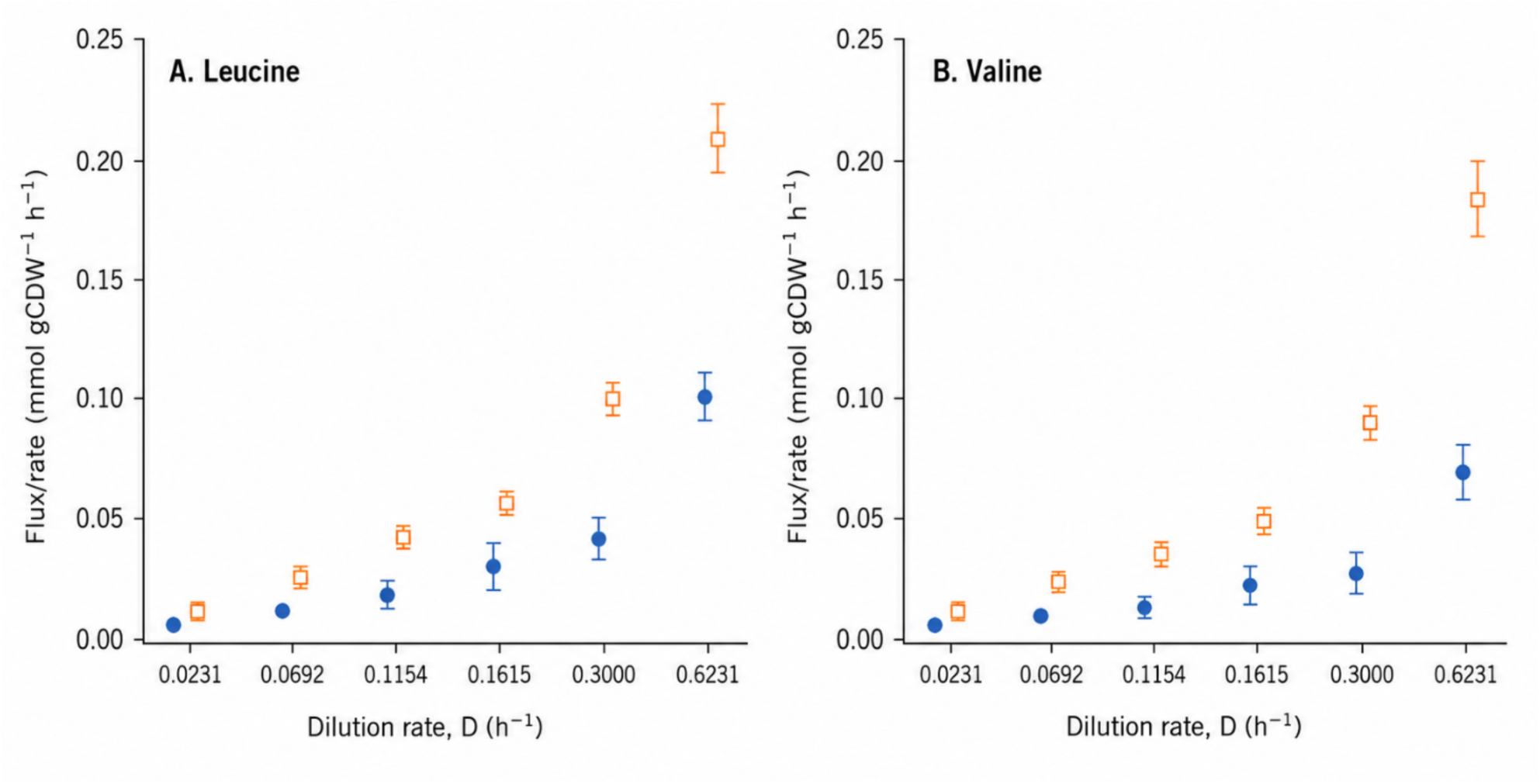
Comparison of leucine and valine rates determined experimentally (blue) with computational model estimates from constrained PMSR7 simulations (orange). (A) Leucine and (B) valine flux/rate comparisons across the interpretable dilution-rate range. Experimental values represent biomass-normalized extracellular amino acid production rates shown as mean ± SD. Computational values represent median leucine- or valine-associated flux estimates from ACHR-sampled constrained PMSR7 solution spaces, with q05–q95 intervals shown where visible. The narrow computational intervals indicate that these reactions were strongly determined under the integrated growth, glucose, oxygen, maintenance, and ¹³C-derived constraints. Experimental amino acid rates were not imposed as model constraints and are shown as independent physiological comparators.

Across the dilution-rate range, the computational estimates were higher than the measured rates, with the largest-condition comparison showing approximately 2.1-fold higher leucine and 2.7-fold higher valine model estimates. This offset is consistent with the fact that the model values represent leucine- and valine-associated intracellular flux requirements under fixed-growth constraints, whereas the experimental values represent cellular accumulation rates. Despite this difference in biological meaning, both datasets showed a clear growth-associated increase and remained within the same order of magnitude. Because cellular leucine and valine measurements were not imposed as model constraints, this comparison supports the physiological plausibility of the integrated PMSR7 framework rather than reflecting direct fitting to the amino acid data.

## Discussion

### PMSR7 provides a quality-benchmarked SR7 reconstruction for constraint-based analysis

PMSR7 establishes a strain-specific genome-scale metabolic model for *Priestia megaterium* SR7 with sufficient structural quality for downstream constraint-based analysis. The reconstruction contains 1,196 genes, 1,424 metabolites, and 1,709 reactions, giving genome-scale coverage comparable to the available *P. megaterium* DSM319 reconstruction while retaining SR7-specific genome association. This provides the biochemical scope needed to examine central carbon metabolism and branched-chain amino acid pathways, which are directly relevant to SR7’s proposed use as a branched-chain alcohol production chassis.

The Memote comparison showed that PMSR7 improved substantially over iJA1121 in structural consistency, increasing overall consistency from 50.69% to 97.15%, mass balance from 83.9% to 99.7%, and charge balance from 82.6% to 91.4%. These improvements are important because mass and charge inconsistencies can create biologically misleading feasible solutions in constraint-based simulations. The remaining annotation gap reflects the challenge of reconstructing metabolism in a newly studied, extremophile-associated *Priestia* strain rather than a failure of model completeness. Compared with canonical model organisms, fewer SR7-specific enzyme functions, transport reactions, and biochemical annotations are experimentally resolved. PMSR7 therefore provides a curated, structurally consistent framework that can incorporate new SR7-specific evidence over time, consistent with the role of genome-scale models as updateable metabolic knowledge bases (Lewis et al., 2012; Thiele & Palsson, 2010).

Together, these results support PMSR7 as a structurally reliable starting model for SR7. The model is not presented as complete because every reaction is experimentally verified, but because it combines genome-scale coverage, improved stoichiometric consistency, and standardized benchmarking. This establishes the foundation required for the subsequent experimental constraint integration using chemostat physiology and ¹³C-MFA-derived flux information.

### Experimental constraint integration makes PMSR7 physiologically usable

The utility of PMSR7 comes not only from reconstruction quality, but from its evaluation under measured *P. megaterium* SR7 physiology. In the chemostat series, each dilution rate defined a steady-state growth condition, allowing biomass flux to be fixed directly to μ = D rather than inferred from unconstrained biomass maximization. This was important because the experimental series spanned a wide growth range, from D = 0.0231 h⁻¹ to D = 0.6231 h⁻¹ for the physiologically interpretable conditions, with the highest dilution rate excluded because biomass collapse made biomass-normalized rates unreliable. Chemostat cultivation therefore provided controlled growth states for model constraint rather than a single endpoint phenotype (Egli, 1995; Monod, 1949; Novick & Szilard, 1950; Pirt, 1975).

Measured glucose uptake provided the main carbon-input constraint. Across the interpretable growth range, biomass-normalized glucose consumption increased from 0.2077 to 5.4851 mmol gCDW⁻¹ h⁻¹, giving condition-specific carbon supply limits for PMSR7. This anchoring was necessary because substrate-unlimited simulations can remain mathematically feasible while permitting physiologically unrealistic carbon throughput. The finite oxygen cap of −25 mmol gCDW⁻¹ h⁻¹ served a similar role by maintaining aerobic metabolism within a bounded solution space rather than allowing unconstrained oxygen uptake. Together with growth-associated ATP demand, these constraints converted the chemostat measurements into explicit model boundaries.

The ¹³C-MFA layer added intracellular information that extracellular measurements alone could not provide. The accepted ¹³C-MFA fit at the representative 0.7 mL min⁻¹ condition had a sum of squared residuals of 127.5, within the expected 95% interval of 92.4–153.3, supporting the use of estimated confidence intervals as intracellular flux constraints. Mapping these bounds into PMSR7 helped restrict central carbon routing beyond what could be inferred from glucose uptake and extracellular amino acid rates alone. Because not all mapped bounds were compatible with the genome-scale model, feasibility screening was required before constraint transfer, which avoided forcing the GEM into infeasible or excessively distorted states. This is consistent with the role of ¹³C-MFA as a complementary intracellular flux measurement framework rather than a replacement for genome-scale stoichiometric analysis (Antoniewicz et al., 2007; Wiechert, 2001; Zamboni, 2011).

The resulting framework is therefore useful because it is explicitly constrained at multiple levels: growth rate, carbon input, oxygen availability, maintenance-energy demand, and intracellular central carbon flux. This does not make PMSR7 a complete predictor of all extracellular products, but it does establish a physiologically bounded model state from which pathway-specific behaviour can be interrogated. For a newly studied, extremophile-associated SR7 chassis, this layered constraint structure is the main advance: it turns PMSR7 from a structural reconstruction into a condition-grounded model that can be refined and extended as additional SR7-specific measurements become available.

### Leucine and valine comparisons support pathway-scale physiological plausibility

The leucine and valine comparison was used as a pathway-scale physiological check rather than as a strict validation of extracellular secretion. This distinction is important because the computational values represent leucine- and valine-associated flux requirements within the constrained model, whereas the experimental values represent biomass-normalized cellular amino acid accumulation.

Within this context, the computational estimates behaved plausibly across the growth series. Leucine-associated model estimates increased from 0.0078 to 0.2096 mmol gCDW⁻¹ h⁻¹, while cellular leucine production increased from 0.0045 ± 0.0006 to 0.1014 ± 0.0035 mmol gCDW⁻¹ h⁻¹. Valine-associated model estimates increased from 0.0069 to 0.1857 mmol gCDW⁻¹ h⁻¹, while cellular valine production increased from 0.0028 ± 0.0003 to 0.0696 ± 0.0015 mmol gCDW⁻¹ h⁻¹. The model estimates were consistently higher than the cellular measurements, particularly at higher dilution rates, but remained within the same order of magnitude and followed the same growth-associated direction.

This offset is biologically reasonable. Leucine and valine are biomass precursors and central intermediates in branched-chain amino acid metabolism, so model-predicted pathway flux is not expected to equal cellular accumulation. The comparison is therefore most informative as an independent scale and trend check: cellular leucine and valine measurements were not imposed as model constraints, yet the constrained PMSR7 framework produced growth-associated flux estimates that increased alongside the measured cellular phenotype. This supports the physiological plausibility of the integrated model without overstating the comparison as direct quantitative prediction.

The narrow ACHR-derived intervals for leucine and valine also provide useful information. Rather than indicating broad feasible-space variability, the near-collapsed intervals suggest that these amino acid-associated fluxes were strongly determined by the fixed-growth formulation and biomass precursor demand under the imposed constraints. This is consistent with the role of leucine and valine as growth-linked biomass components and reinforces why these values are best interpreted as growth-coupled computational flux estimates, not as freely varying secretion predictions. For SR7 engineering, this result is still valuable because branched-chain amino acid metabolism supplies keto-acid precursors for branched alcohol production, including isobutanol and isopentanol pathways (Atsumi et al., 2008; Boock et al., 2019; Connor & Liao, 2008).

### PMSR7 as a platform for *P. megaterium* SR7 metabolic engineering

The main practical value of PMSR7 is that it provides a condition-grounded baseline for testing *P. megaterium* SR7 metabolic engineering hypotheses. Because the model is constrained by measured growth, glucose uptake, oxygen availability, maintenance-energy demand, and intracellular ¹³C-MFA information, it can be used to examine how carbon is allocated through central metabolism under defined physiological states. This is especially relevant for *P. megaterium* SR7 engineering because branched-chain alcohol production depends on precursor supply through valine- and leucine-associated keto-acid pathways, rather than on the final alcohol pathway alone (Atsumi et al., 2008; Boock et al., 2019; Connor & Liao, 2008).

In this form, PMSR7 can support future design questions such as whether growth state favours precursor availability, which reactions constrain flux toward 2-ketoisovalerate- or leucine-derived intermediates, and how oxygen or substrate constraints affect the balance between biomass formation and production-associated carbon withdrawal. These uses are more appropriate than treating the model as a direct predictor of extracellular product titres, particularly because the present framework was developed under aerobic, glucose-limited wild-type chemostat conditions. Additional constraints would be needed before applying the model quantitatively to engineered strains, CO₂-enriched cultivation, solvent-extractive systems, or alternative carbon sources.

PMSR7 also provides an updateable framework for incorporating future *P. megaterium* SR7-specific datasets. Additional isotope-labelling experiments across selected dilution rates could test whether central carbon routing changes with growth rate, while engineered-strain measurements could be used to constrain heterologous branched alcohol or PHB production pathways. As more *P. megaterium* SR7-specific enzyme annotations, transport functions, and production phenotypes become available, PMSR7 can be refined from a growth-resolved wild-type baseline into a predictive platform for chassis development. Thus, the model introduced here is most useful as a structured, experimentally anchored starting point for iterative *P. megaterium* SR7 metabolic design.

## Conclusion

This study presents PMSR7, a genome-scale metabolic model for *P. megaterium* SR7 and demonstrates its use under experimentally defined aerobic chemostat conditions. PMSR7 showed improved structural consistency relative to the available *P. megaterium* reconstruction and provided genome-scale coverage suitable for analysing central carbon and branched-chain amino acid metabolism. By integrating dilution-rate-defined growth, measured glucose uptake, bounded oxygen availability, growth-associated ATP demand, and ¹³C-MFA-derived intracellular flux constraints, the model was converted from a structural reconstruction into a condition-grounded framework for *P. megaterium* SR7 metabolic interpretation.

The constrained model produced growth-associated leucine and valine flux estimates that were consistent in scale and direction with independently measured cellular amino acid rates. These results support PMSR7 as a usable baseline model for interrogating SR7 metabolism and for guiding future metabolic engineering hypotheses involving branched-chain amino acid-derived products. As additional *P. megaterium* SR7-specific biochemical, isotope-labelling, and engineered-strain datasets become available, PMSR7 can be iteratively refined into a predictive platform for chassis development under production-relevant conditions.

## Acknowledgements

This research was supported by the Singapore Ministry of Education under Academic Research Fund Tier 2 (MOE-T2EP30122-0015) and by the Singapore Ministry of Education and National Research Foundation through a RCE award to Singapore Centre for Environmental Life Sciences Engineering (SCELSE).

## Supplementary

### S1 C13-MFA

**S1.1.**
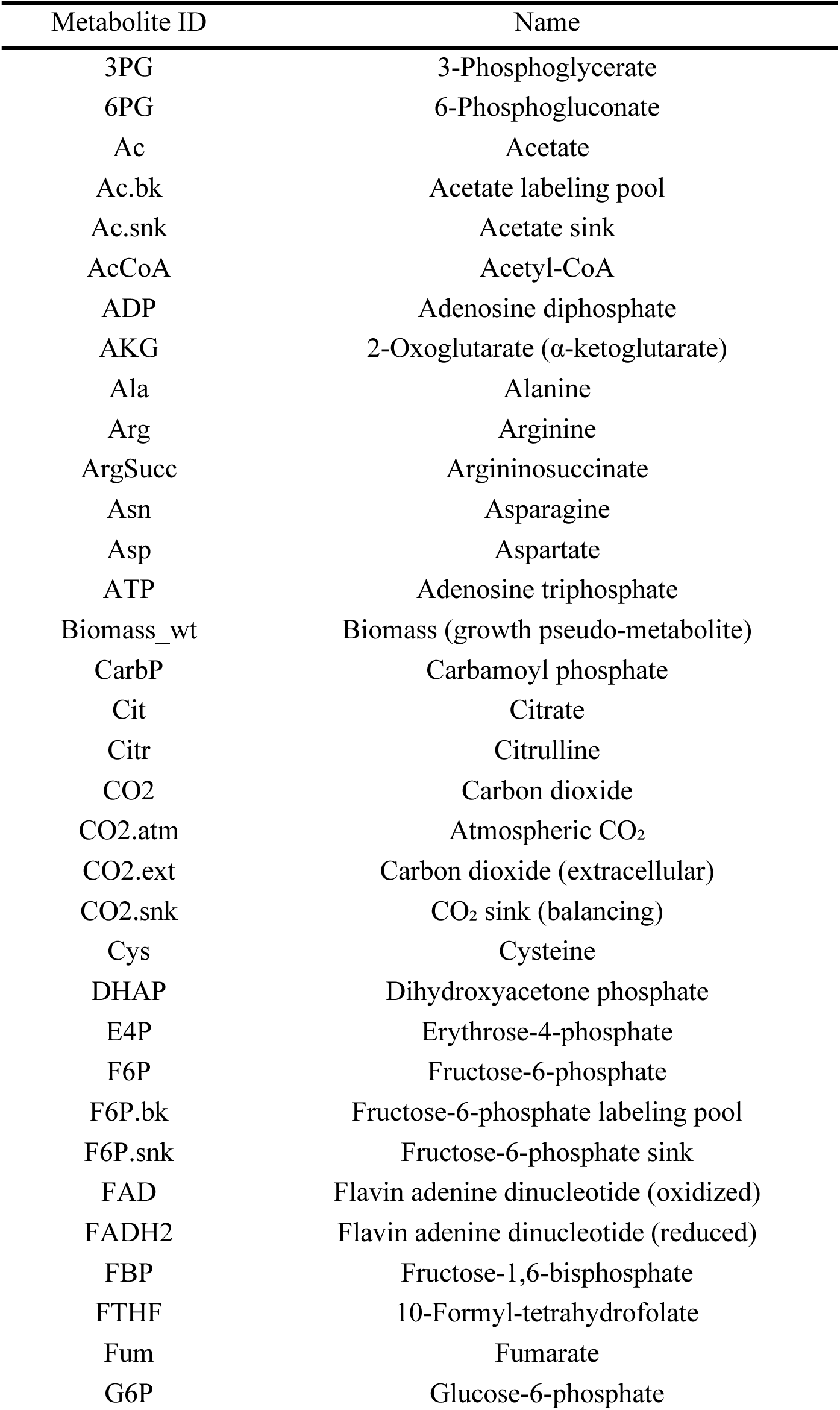
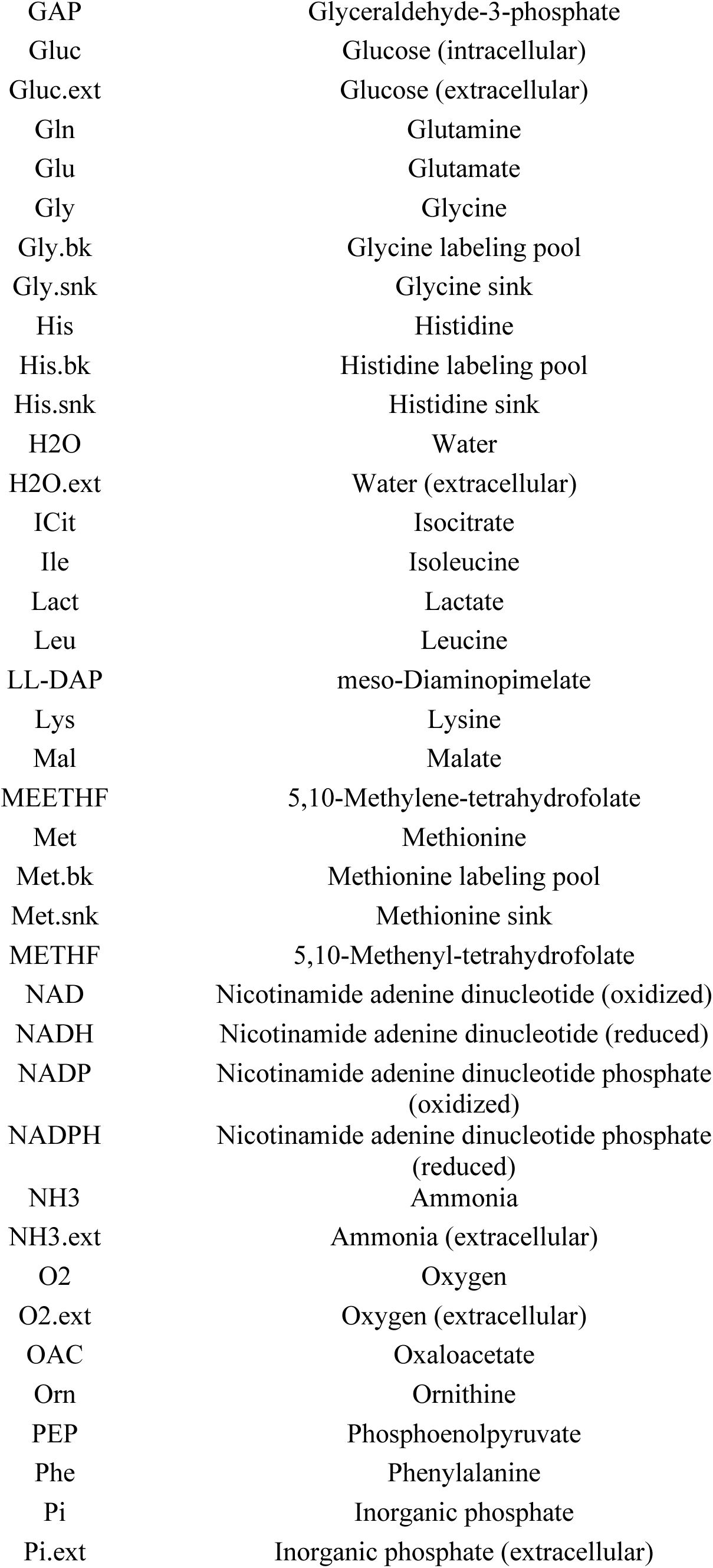
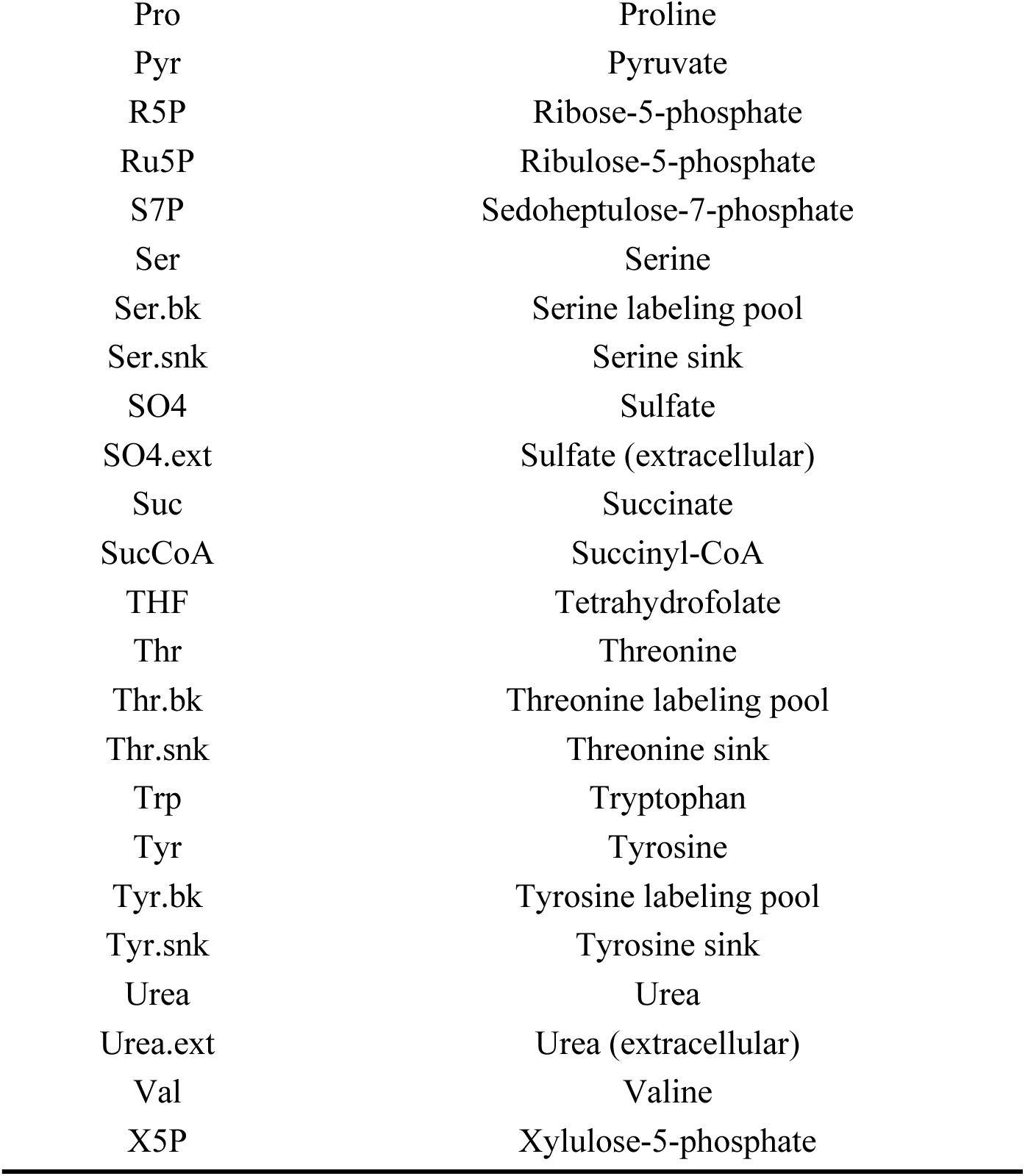
Showing MFA model metabolite IDs

S1.1 Showing MFA model metabolite IDs
| Metabolite ID | Name |
| --- | --- |
| 3PG | 3-Phosphoglycerate |
| 6PG | 6-Phosphogluconate |
| Ac | Acetate |
| Ac.bk | Acetate labeling pool |
| Ac.snk | Acetate sink |
| AcCoA | Acetyl-CoA |
| ADP | Adenosine diphosphate |
| AKG | 2-Oxoglutarate (α-ketoglutarate) |
| Ala | Alanine |
| Arg | Arginine |
| ArgSucc | Argininosuccinate |
| Asn | Asparagine |
| Asp | Aspartate |
| ATP | Adenosine triphosphate |
| Biomass_wt | Biomass (growth pseudo-metabolite) |
| CarbP | Carbamoyl phosphate |
| Cit | Citrate |
| Citr | Citrulline |
| CO2 | Carbon dioxide |
| CO2.atm | Atmospheric CO<sub>2</sub> |
| CO2.ext | Carbon dioxide (extracellular) |
| CO2.snk | CO<sub>2</sub> sink (balancing) |
| Cys | Cysteine |
| DHAP | Dihydroxyacetone phosphate |
| E4P | Erythrose-4-phosphate |
| F6P | Fructose-6-phosphate |
| F6P.bk | Fructose-6-phosphate labeling pool |
| F6P.snk | Fructose-6-phosphate sink |
| FAD | Flavin adenine dinucleotide (oxidized) |
| FADH2 | Flavin adenine dinucleotide (reduced) |
| FBP | Fructose-1,6-bisphosphate |
| FTHF | 10-Formyl-tetrahydrofolate |
| Fum | Fumarate |
| G6P | Glucose-6-phosphate |
| GAP | Glyceraldehyde-3-phosphate |
| Gluc | Glucose (intracellular) |
| Gluc.ext | Glucose (extracellular) |
| Gln | Glutamine |
| Glu | Glutamate |
| Gly | Glycine |
| Gly.bk | Glycine labeling pool |
| Gly.snk | Glycine sink |
| His | Histidine |
| His.bk | Histidine labeling pool |
| His.snk | Histidine sink |
| H2O | Water |
| H2O.ext | Water (extracellular) |
| ICit | Isocitrate |
| Ile | Isoleucine |
| Lact | Lactate |
| Leu | Leucine |
| LL-DAP | meso-Diaminopimelate |
| Lys | Lysine |
| Mal | Malate |
| MEETHF | 5,10-Methylene-tetrahydrofolate |
| Met | Methionine |
| Met.bk | Methionine labeling pool |
| Met.snk | Methionine sink |
| METHF | 5,10-Methenyl-tetrahydrofolate |
| NAD | Nicotinamide adenine dinucleotide (oxidized) |
| NADH | Nicotinamide adenine dinucleotide (reduced) |
| NADP | Nicotinamide adenine dinucleotide phosphate (oxidized) |
| NADPH | Nicotinamide adenine dinucleotide phosphate (reduced) |
| NH3 | Ammonia |
| NH3.ext | Ammonia (extracellular) |
| O2 | Oxygen |
| O2.ext | Oxygen (extracellular) |
| OAC | Oxaloacetate |
| Orn | Ornithine |
| PEP | Phosphoenolpyruvate |
| Phe | Phenylalanine |
| Pi | Inorganic phosphate |
| Pi.ext | Inorganic phosphate (extracellular) |
| Pro | Proline |
| Pyr | Pyruvate |
| R5P | Ribose-5-phosphate |
| Ru5P | Ribulose-5-phosphate |
| S7P | Sedoheptulose-7-phosphate |
| Ser | Serine |
| Ser.bk | Serine labeling pool |
| Ser.snk | Serine sink |
| SO4 | Sulfate |
| SO4.ext | Sulfate (extracellular) |
| Suc | Succinate |
| SucCoA | Succinyl-CoA |
| THF | Tetrahydrofolate |
| Thr | Threonine |
| Thr.bk | Threonine labeling pool |
| Thr.snk | Threonine sink |
| Trp | Tryptophan |
| Tyr | Tyrosine |
| Tyr.bk | Tyrosine labeling pool |
| Tyr.snk | Tyrosine sink |
| Urea | Urea |
| Urea.ext | Urea (extracellular) |
| Val | Valine |
| X5P | Xylulose-5-phosphate |

**S1.2.** Showing lumped output reactions and C13-MFA confidence intervals. The C13-MFA solution was normalized such that the glucose uptake reaction (Gluc.ext + PEP -> G6P + Pyr) was set to 100. Scaled fluxes were then rescaled to the experimental glucose uptake at 0.7 ml/min (1.6165 mmol·gCDW⁻¹·h⁻¹).

| No | Reaction | 95% CI | Scaled 95% CI |
| --- | --- | --- | --- |
| 1 | $\text{Gluc.ext} + \text{PEP} \rightarrow \text{G6P} + \text{Pyr}$ | ( 41.53, 158.48 ) | ( 0.67, 2.56 ) |
| 2 | $\text{G6P} \rightarrow \text{F6P}$ | ( 5.13, 87.65 ) | ( 0.08, 1.42 ) |
| 3 | $\text{F6P} + \text{ATP} \rightarrow \text{FBP} + \text{ADP}$ | ( 50.72, 97.70 ) | ( 0.82, 1.58 ) |
| 4 | $\text{FBP} \rightleftharpoons \text{DHAP} + \text{GAP} \text{ (net)}$ | ( 50.72, 97.70 ) | ( 0.82, 1.58 ) |
| 5 | $\text{FBP} \rightleftharpoons \text{DHAP} + \text{GAP} \text{ (exch)}$ | ( 0.00, >10000 ) | ( 0.00, 161.49 ) |
| 6 | $\text{DHAP} \rightleftharpoons \text{GAP} \text{ (net)}$ | ( 50.72, 97.70 ) | ( 0.82, 1.58 ) |
| 7 | $\text{DHAP} \rightleftharpoons \text{GAP} \text{ (exch)}$ | ( 0.00, >10000 ) | ( 0.00, 161.49 ) |
| 8 | $\text{GAP} + \text{NAD} + \text{ADP} + \text{Pi} \rightleftharpoons \text{3PG} + \text{ATP} + \text{NADH} \text{ (net)}$ | ( 129.84, 200.71 ) | ( 2.10, 3.24 ) |
| 9 | $\text{GAP} + \text{NAD} + \text{ADP} + \text{Pi} \rightleftharpoons \text{3PG} + \text{ATP} + \text{NADH} \text{ (exch)}$ | ( 0.00, >10000 ) | ( 0.00, 161.49 ) |
| 10 | $\text{3PG} \rightleftharpoons \text{PEP} \text{ (net)}$ | ( 132.60, 200.23 ) | ( 2.14, 3.23 ) |
| 11 | 3PG <=> PEP (exch) | ( 0.00, >10000 ) | ( 0.00, 161.49 ) |
| 12 | PEP + ADP <=> Pyr + ATP (net) | ( -100.00, 94.79 ) | ( -1.61, 1.53 ) |
| 13 | PEP + ADP <=> Pyr + ATP (exch) | ( 0.00, >10000 ) | ( 0.00, 161.49 ) |
| 14 | G6P + NADP -> 6PG + NADPH | ( 9.47, 216.83 ) | ( 0.15, 3.50 ) |
| 15 | 6PG + NADP -> Ru5P + CO2 + NADPH | ( 9.47, 216.83 ) | ( 0.15, 3.50 ) |
| 16 | Ru5P <=> X5P (net) | ( 6.09, 10.65 ) | ( 0.10, 0.17 ) |
| 17 | Ru5P <=> X5P (exch) | ( 0.00, 3.14 ) | ( 0.00, 0.05 ) |
| 18 | Ru5P <=> R5P (net) | ( 2.00, 71.22 ) | ( 0.03, 1.15 ) |
| 19 | Ru5P <=> R5P (exch) | ( 0.00, 0.99 ) | ( 0.00, 0.02 ) |
| 20 | Pyr + NAD -> AcCoA + CO2 + NADH | ( 0.01, 15.64 ) | ( 0.00, 0.25 ) |
| 21 | AcCoA + OAC -> Cit | ( 0.01, 6.31 ) | ( 0.00, 0.10 ) |
| 22 | Cit <=> ICit (net) | ( 0.01, 6.31 ) | ( 0.00, 0.10 ) |
| 23 | Cit <=> ICit (exch) | ( 0.00, >10000 ) | ( 0.00, 161.49 ) |
| 24 | ICit + NADP <=> AKG + CO2 + NADPH (net) | ( 0.01, 6.31 ) | ( 0.00, 0.10 ) |
| 25 | ICit + NADP <=> AKG + CO2 + NADPH (exch) | ( 0.00, >10000 ) | ( 0.00, 161.49 ) |
| 26 | AKG + NAD <=> SuccCoA + CO2 + NADH (net) | ( 0.00, 8.48 ) | ( 0.00, 0.14 ) |
| 27 | AKG + NAD <=> SuccCoA + CO2 + NADH (exch) | ( 3.67, >10000 ) | ( 0.06, 161.49 ) |
| 28 | SuccCoA + ADP + Pi <=> Suc + ATP (net) | ( -0.75, -0.00 ) | ( -0.01, -0.00 ) |
| 29 | SuccCoA + ADP + Pi <=> Suc + ATP (exch) | ( 0.01, 1.10 ) | ( 0.00, 0.02 ) |
| 30 | Suc + FAD <=> Fum + FADH2 (net) | ( 0.00, 8.48 ) | ( 0.00, 0.14 ) |
| 31 | Suc + FAD <=> Fum + FADH2 (exch) | ( 8.38, >10000 ) | ( 0.14, 161.49 ) |
| 32 | Fum <=> Mal (net) | ( 0.00, 10.21 ) | ( 0.00, 0.16 ) |
| 33 | Fum <=> Mal (exch) | ( 0.00, >10000 ) | ( 0.00, 161.49 ) |
| 34 | Mal + NAD <=> OAC + NADH (net) | ( -251.73, 31.38 ) | ( -4.06, 0.51 ) |
| 35 | Mal + NAD <=> OAC + NADH (exch) | ( 0.00, 550.71 ) | ( 0.00, 8.89 ) |
| 36 | Mal + NADP -> Pyr + CO2 + NADPH | ( 0.06, 444.54 ) | ( 0.00, 7.18 ) |
| 37 | PEP + CO2 -> OAC + Pi | ( 0.11, 452.13 ) | ( 0.00, 7.30 ) |
| 38 | AcCoA + ADP + Pi -> Ac + ATP | ( 0.00, 172.27 ) | ( 0.00, 2.78 ) |
| 39 | AKG + NADPH + NH3 -> Glu + NADP | ( 0.07, 38.59 ) | ( 0.00, 0.62 ) |
| 40 | Glu + ATP + NH3 -> Gln + ADP + Pi | ( 0.01, 5.95 ) | ( 0.00, 0.10 ) |
| 41 | Glu + 2 NADPH + ATP -> Pro + 2 NADP + ADP + Pi | ( 0.00, 0.49 ) | ( 0.00, 0.01 ) |
| 42 | Glu + CO2 + Gln + NADPH + Asp + AcCoA + 5 ATP -> Arg + AKG + NADP + Fum + Ac + 5 ADP + 5 Pi | ( 0.00, 0.30 ) | ( 0.00, 0.00 ) |
| 43 | OAC + Glu -> Asp + AKG | ( 0.02, 12.30 ) | ( 0.00, 0.20 ) |
| 44 | Asp + NH3 + 2 ATP -> Asn + 2 ADP + 2 Pi | ( 0.00, 0.40 ) | ( 0.00, 0.01 ) |
| 45 | 3PG + Glu + NAD -> Ser + NADH + AKG + Pi | ( 0.00, 0.82 ) | ( 0.00, 0.01 ) |
| 46 | Ser + THF -> Gly + METHF | ( 0.00, 0.28 ) | ( 0.00, 0.00 ) |
| 47 | Gly + THF + NAD <=> CO2 + MEETHF + NH3 + NADH (net) | ( 0.00, 0.33 ) | ( 0.00, 0.01 ) |
| 48 | Gly + THF + NAD <=> CO2 + MEETHF + NH3 + NADH (exch) | ( 0.00, 0.24 ) | ( 0.00, 0.00 ) |
| 49 | Thr + NAD -> Gly + AcCoA + NADH | ( 0.00, 1.71 ) | ( 0.00, 0.03 ) |
| 50 | Ser + AcCoA + SO4 + 3 ATP + 4 NADPH -> Cys + Ac + 4 NADP + 3 ADP + 3 Pi | ( 0.00, 0.36 ) | ( 0.00, 0.01 ) |
| 51 | Asp + Pyr + Glu + 2 NADPH + ATP + SucCoA -> LL-DAP + AKG + 2 NADP + ADP + Pi + Suc | ( 0.00, 0.56 ) | ( 0.00, 0.01 ) |
| 52 | LL-DAP -> Lys + CO2 | ( 0.00, 0.56 ) | ( 0.00, 0.01 ) |
| 53 | Asp + 2 NADPH + 2 ATP -> Thr + 2 NADP + 2 ADP + 2 Pi | ( 0.00, 2.68 ) | ( 0.00, 0.04 ) |
| 54 | Asp + METHF + Cys + 2 NADPH + ATP + SucCoA -> Met + Pyr + 2 NADP + ADP + Pi + Suc + NH3 + THF | ( 0.00, 0.19 ) | ( 0.00, 0.00 ) |
| 55 | 2 Pyr + NADPH + Glu -> Val + CO2 + NADP + AKG | ( 0.00, 0.44 ) | ( 0.00, 0.01 ) |
| 56 | 2 Pyr + AcCoA + Glu + NADPH + NAD -> Leu + 2 CO2 + AKG + NADP + NADH | ( 0.00, 0.71 ) | ( 0.00, 0.01 ) |
| 57 | Thr + Pyr + Glu + NADPH -> Ile + CO2 + AKG + NADP + NH3 | ( 0.00, 0.42 ) | ( 0.00, 0.01 ) |
| 58 | E4P + 2 PEP + Glu + NADPH + ATP -> Phe + CO2 + AKG + NADP + ADP + Pi | ( 0.00, 0.31 ) | ( 0.00, 0.01 ) |
| 59 | E4P + 2 PEP + Glu + NADPH + NAD + ATP -> Tyr + CO2 + AKG + NADP + ADP + Pi | ( 0.00, 0.18 ) | ( 0.00, 0.00 ) |
| 60 | E4P + 2 PEP + R5P + Ser + Gln + NADPH + 3 ATP -> Trp + CO2 + 3 ADP + 3 Pi + NADP | ( 0.00, 0.01 ) | ( 0.00, 0.00 ) |
| 61 | R5P + FTHF + Gln + Asp + 5 ATP + 2 NAD -> His + 2 NADH + 5 ADP + 5 Pi | ( 0.00, 0.14 ) | ( 0.00, 0.00 ) |
| 62 | MEETHF + NADH -> METHF + NAD | ( 0.00, 0.19 ) | ( 0.00, 0.00 ) |
| 63 | MEETHF + NADP -> FTHF + NADPH | ( 0.00, 0.14 ) | ( 0.00, 0.00 ) |
| 64 | NADH + 0.5 O2 + 1.5 ADP + 1.5 Pi -> NAD + 1.5 ATP | ( 12.23, 98.50 ) | ( 0.20, 1.59 ) |
| 65 | FADH2 + 0.5 O2 + 0.5 ADP + 0.5 Pi -> FAD + 0.5 ATP | ( 0.00, 8.48 ) | ( 0.00, 0.14 ) |
| 66 | NADH + NADP <=> NADPH + NAD (net) | ( -31.74, -14.49 ) | ( -0.51, -0.23 ) |
| 67 | NADH + NADP <=> NADPH + NAD (exch) | ( 0.00, >10000 ) | ( 0.00, 161.49 ) |
| 68 | ATP -> ADP + Pi | ( 192.30, 240.76 ) | ( 3.11, 3.89 ) |
| 69 | Ac -> Ac.ext | ( 0.00, 173.72 ) | ( 0.00, 2.80 ) |
| 70 | CO2 -> CO2.ext | ( 8.28, 265.35 ) | ( 0.13, 4.28 ) |
| 71 | O2.ext -> O2 | ( 6.12, 63.08 ) | ( 0.10, 1.02 ) |
| 72 | NH3.ext -> NH3 | ( 0.09, 70.17 ) | ( 0.00, 1.13 ) |
| 73 | SO4.ext -> SO4 | ( 0.00, 0.36 ) | ( 0.00, 0.01 ) |
| 74 | Pi.ext -> Pi | ( 0.01, 5.65 ) | ( 0.00, 0.09 ) |
| 75 | CO2.atm + CO2 -> CO2 + CO2.snk | ( 12.97, 22.98 ) | ( 0.21, 0.37 ) |
| 76 | Pyr + NADH -> Lact + NAD (net) | ( 141.40, 195.20 ) | ( 2.28, 3.15 ) |
| 77 | Pyr + NADH <=> Lact + NAD (exch) | ( 0.00, >10000 ) | ( 0.00, 161.49 ) |
| 78 | Ala <=> Pyr (net) | ( -1.64, -0.00 ) | ( -0.03, -0.00 ) |
| 79 | Ala <=> Pyr (exch) | ( 0.00, >10000 ) | ( 0.00, 161.49 ) |
| 80 | Lact -> Lact.ext | ( 141.40, 195.20 ) | ( 2.28, 3.15 ) |
| 81 | Ser.bk <=> Ser (net) | ( 0.00, 0.00 ) | ( 0.00, 0.00 ) |
| 82 | Ser.bk <=> Ser (exch) | ( 0.00, >10000 ) | ( 0.00, 161.49 ) |
| 83 | Ser.bk <=> Ser.snk (net) | ( 0.00, 0.00 ) | ( 0.00, 0.00 ) |
| 84 | Ser.bk <=> Ser.snk (exch) | ( 0.00, >10000 ) | ( 0.00, 161.49 ) |
| 85 | Gly.bk <=> Gly (net) | ( 0.00, 0.00 ) | ( 0.00, 0.00 ) |
| 86 | Gly.bk <=> Gly (exch) | ( 0.00, >10000 ) | ( 0.00, 161.49 ) |
| 87 | Gly.bk <=> Gly.snk (net) | ( 0.00, 0.00 ) | ( 0.00, 0.00 ) |
| 88 | Gly.bk <=> Gly.snk (exch) | ( 0.00, >10000 ) | ( 0.00, 161.49 ) |
| 89 | His.bk <=> His (net) | ( 0.00, 0.00 ) | ( 0.00, 0.00 ) |
| 90 | His.bk <=> His (exch) | ( 0.00, >10000 ) | ( 0.00, 161.49 ) |
| 91 | His.bk <=> His.snk (net) | ( 0.00, 0.00 ) | ( 0.00, 0.00 ) |
| 92 | His.bk <=> His.snk (exch) | ( 0.00, >10000 ) | ( 0.00, 161.49 ) |
| 93 | F6P.bk <=> F6P (net) | ( 0.00, 0.00 ) | ( 0.00, 0.00 ) |
| 94 | F6P.bk <=> F6P (exch) | ( 0.00, >10000 ) | ( 0.00, 161.49 ) |
| 95 | F6P.bk <=> F6P.snk (net) | ( 0.00, 0.00 ) | ( 0.00, 0.00 ) |
| 96 | F6P.bk <=> F6P.snk (exch) | ( 0.00, >10000 ) | ( 0.00, 161.49 ) |
| 97 | Tyr.bk <=> Tyr (net) | ( 0.00, 0.00 ) | ( 0.00, 0.00 ) |
| 98 | Tyr.bk <=> Tyr (exch) | ( 0.00, >10000 ) | ( 0.00, 161.49 ) |
| 99 | Tyr.bk <=> Tyr.snk (net) | ( 0.00, 0.00 ) | ( 0.00, 0.00 ) |
| 100 | Tyr.bk <=> Tyr.snk (exch) | ( 0.00, >10000 ) | ( 0.00, 161.49 ) |
| 101 | Thr.bk <=> Thr (net) | ( 0.00, 0.00 ) | ( 0.00, 0.00 ) |
| 102 | Thr.bk <=> Thr (exch) | ( 0.00, >10000 ) | ( 0.00, 161.49 ) |
| 103 | Thr.bk <=> Thr.snk (net) | ( 0.00, 0.00 ) | ( 0.00, 0.00 ) |
| 104 | Thr.bk <=> Thr.snk (exch) | ( 0.00, >10000 ) | ( 0.00, 161.49 ) |
| 105 | 0.42584 Ala + 0.07719 Arg + 0.10238 Asn + 0.21233<br>Asp + 0.042673 Cys + 0.14248 Gln + 0.35688 Gly +<br>0.036846 His + 0.10793 Ile + 0.18403 Leu + 0.14615<br>Lys + 0.049279 Met + 0.079672 Phe + 0.12618 Pro +<br>0.10968 Ser + 0.14176 Thr + 0.0026671 Trp + 0.045586<br>Tyr + 0.11317 Val + 0.25508 G6P + 0.7147 R5P +<br>0.44235 GAP + 0.050756 PEP + 1.2048 AcCoA -><br>Biomass_wt | ( 0.00, 3.86 ) | ( 0.00, 0.06 ) |
| 106 | X5P + R5P <=> GAP + S7P (net) | ( 3.03, 5.37 ) | ( 0.05, 0.09 ) |
| 107 | X5P + R5P <=> GAP + S7P (exch) | ( 3.99, 61.79 ) | ( 0.06, 1.00 ) |
| 108 | X5P + E4P <=> GAP + F6P (net) | ( 3.10, 5.30 ) | ( 0.05, 0.09 ) |
| 109 | X5P + E4P <=> GAP + F6P (exch) | ( 0.00, 3.77 ) | ( 0.00, 0.06 ) |
| 110 | S7P + GAP <=> F6P + E4P (net) | ( 3.03, 5.37 ) | ( 0.05, 0.09 ) |
| 111 | S7P + GAP <=> F6P + E4P (exch) | ( 10.31, >10000<br>) | ( 0.17, 161.49 ) |
| 112 | Ac.bk <=> Ac (net) | ( 0.00, 0.00 ) | ( 0.00, 0.00 ) |
| 113 | Ac.bk <=> Ac (exch) | ( 0.00, >10000 ) | ( 0.00, 161.49 ) |
| 114 | Ac.bk <=> Ac.snk (net) | ( 0.00, 0.00 ) | ( 0.00, 0.00 ) |
| 115 | Ac.bk <=> Ac.snk (exch) | ( 0.00, >10000 ) | ( 0.00, 161.49 ) |
| 116 | Met.bk <=> Met (net) | ( 0.00, 0.00 ) | ( 0.00, 0.00 ) |
| 117 | Met.bk <=> Met (exch) | ( 0.00, >10000 ) | ( 0.00, 161.49 ) |
| 118 | Met.bk <=> Met.snk (net) | ( 0.00, 0.00 ) | ( 0.00, 0.00 ) |
| 119 | Met.bk <=> Met.snk (exch) | ( 0.00, >10000 ) | ( 0.00, 161.49 ) |
| 120 | CO2 + NH3 + 2 ATP -> CarbP + 2 ADP + Pi | ( 0.00, 10.16 ) | ( 0.00, 0.16 ) |
| 121 | CarbP + Orn -> Citr + Pi | ( 0.00, 10.16 ) | ( 0.00, 0.16 ) |
| 122 | Citr + Asp + ATP -> ArgSucc + ADP + Pi | ( 0.00, 10.16 ) | ( 0.00, 0.16 ) |
| 123 | ArgSucc -> Arg + Fum | ( 0.00, 10.16 ) | ( 0.00, 0.16 ) |
| 124 | Arg + H2O -> Orn + Urea | ( 0.00, 10.16 ) | ( 0.00, 0.16 ) |
| 125 | CO2.ext -> CO2 | ( 12.97, 22.98 ) | ( 0.21, 0.37 ) |
| 126 | Gluc.ext -> Gluc | ( 41.53, 158.48 ) | ( 0.67, 2.56 ) |
| 127 | O2.ext -> O2 | ( 6.12, 63.08 ) | ( 0.10, 1.02 ) |
| 128 | NH3.ext -> NH3 | ( 0.09, 70.17 ) | ( 0.00, 1.13 ) |
| 129 | H2O.ext -> H2O | ( 0.00, 10.16 ) | ( 0.00, 0.16 ) |
| 130 | Pi.ext -> Pi | ( 0.01, 5.65 ) | ( 0.00, 0.09 ) |
| 131 | SO4.ext -> SO4 | ( 0.00, 0.36 ) | ( 0.00, 0.01 ) |
| 132 | Ac -> Ac.ext | ( 0.00, 173.72 ) | ( 0.00, 2.80 ) |
| 133 | CO2 -> CO2.ext | ( 8.28, 265.35 ) | ( 0.13, 4.28 ) |
| 134 | CO2 -> CO2.snk | ( 12.97, 22.98 ) | ( 0.21, 0.37 ) |
| 135 | Lact -> Lact.ext | ( 141.40, 195.20 ) | ( 2.28, 3.15 ) |
| 136 | Biomass -> Biomass_wt | ( 0.00, 3.86 ) | ( 0.00, 0.06 ) |
| 137 | Urea -> Urea.ext | ( 0.00, 10.16 ) | ( 0.00, 0.16 ) |

**Appendix S1.3.**
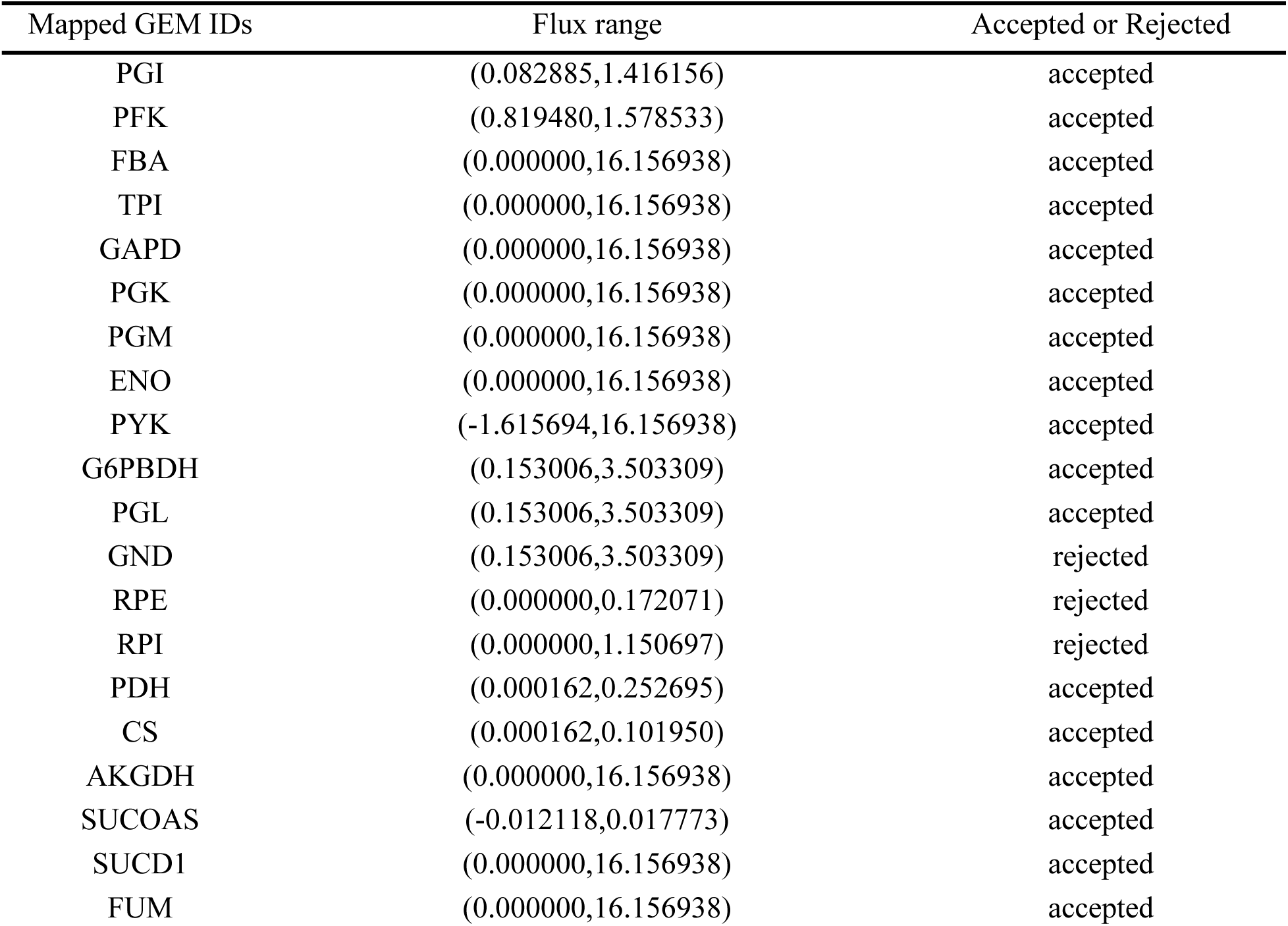
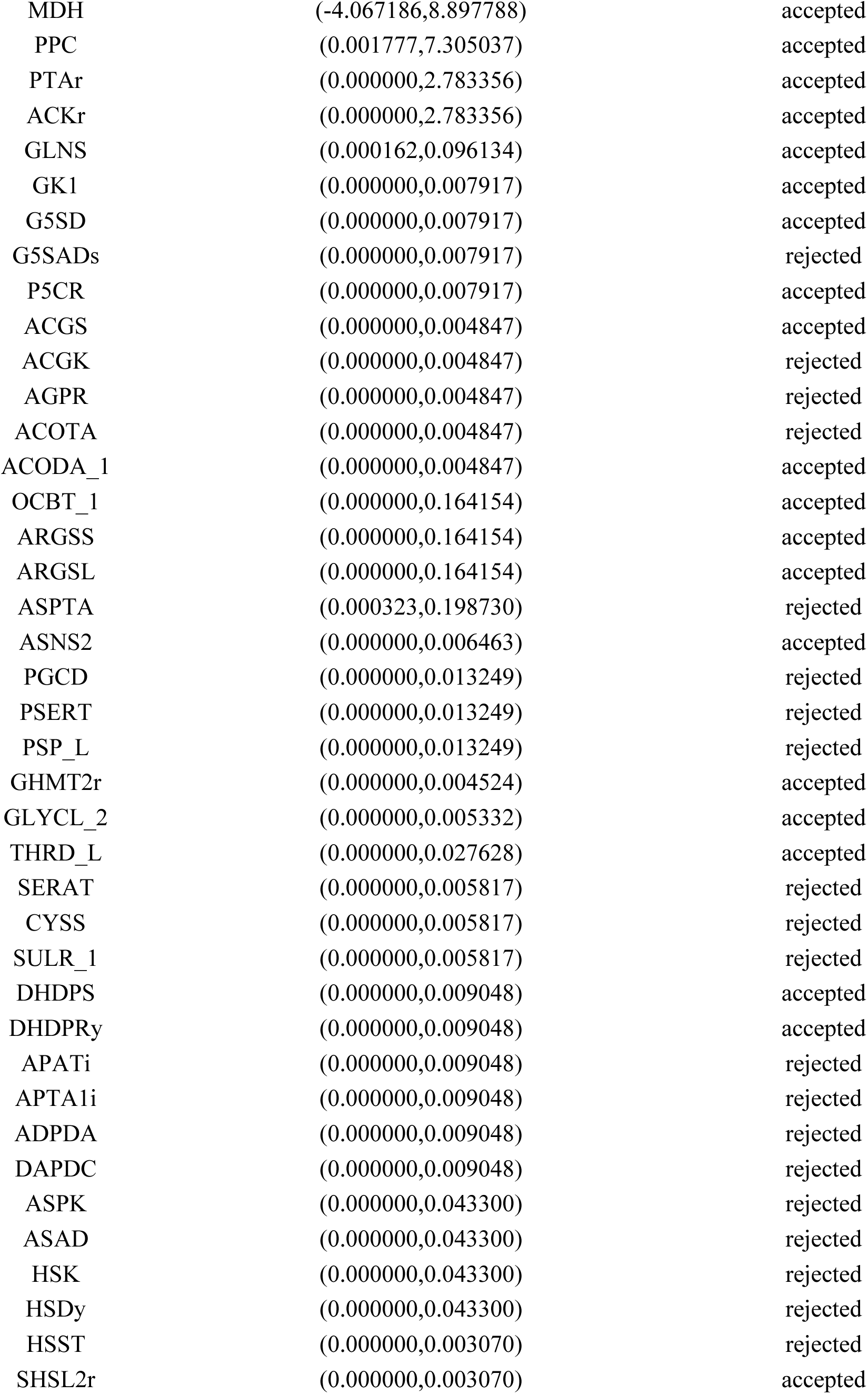
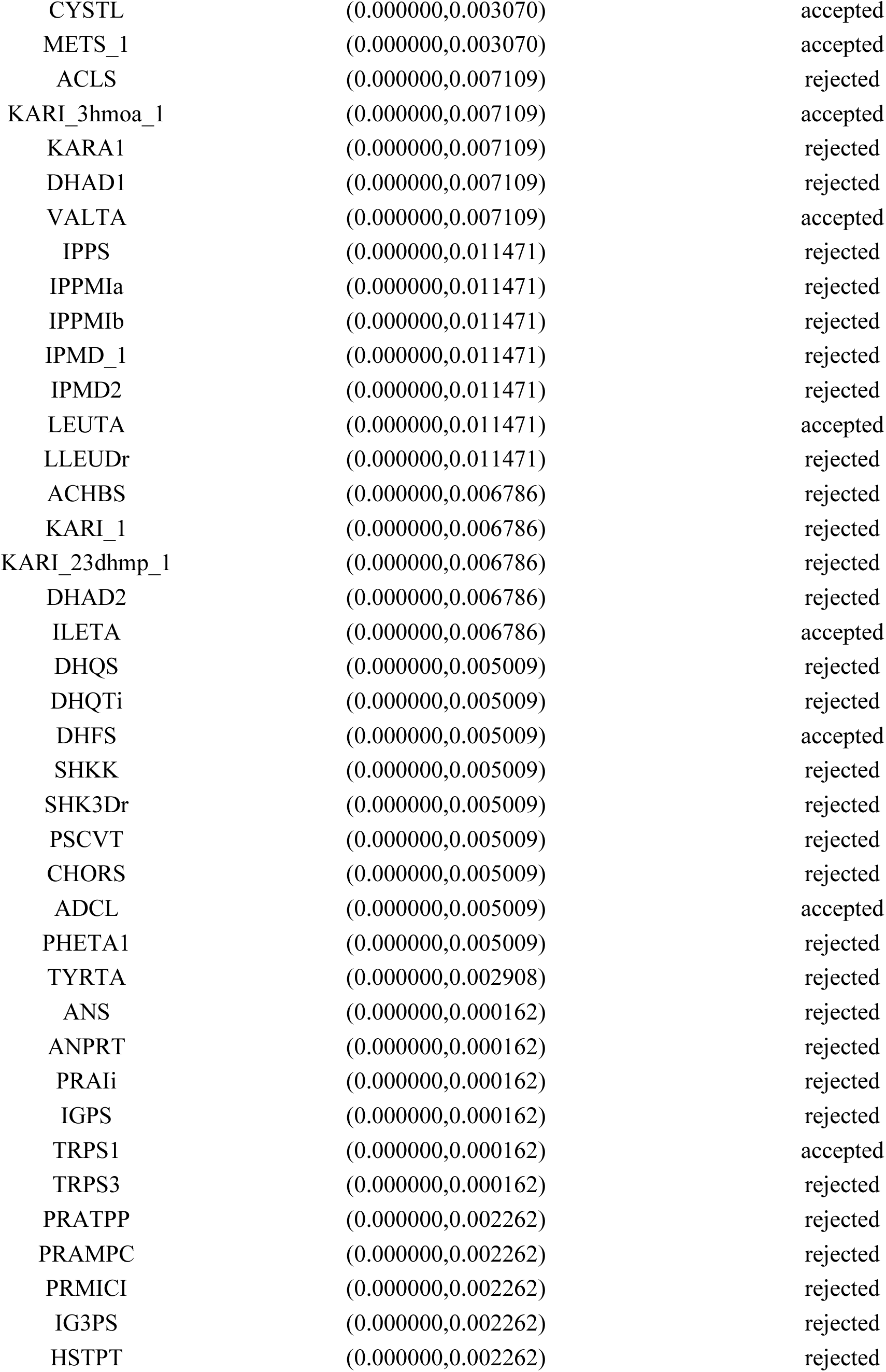
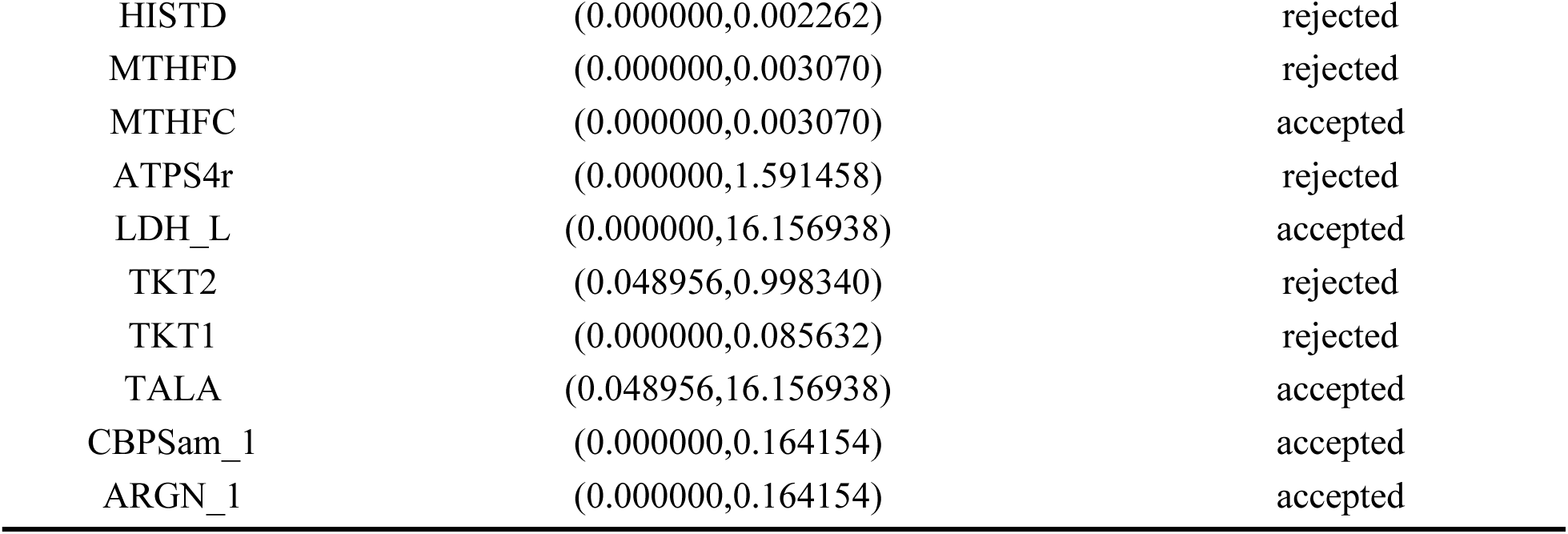
Showing mapped GEM IDs with C13-MFA reaction bounds that were applied (accepted) to the GEM or were left unusued (rejected) due to infeasibility.

### S2 ATP for GAM

**S2.1.**
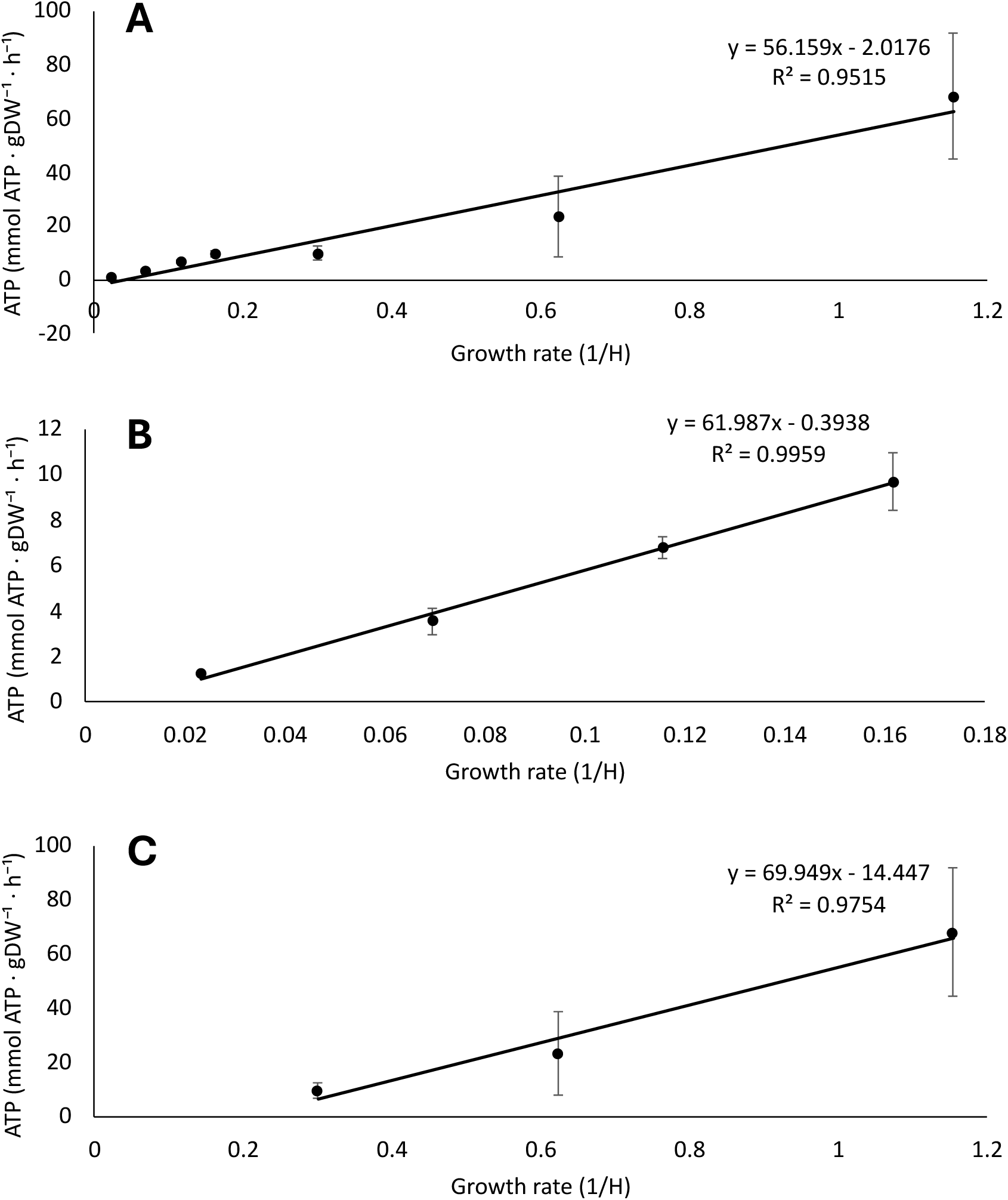
Showing modelled ATP maintenance against growth rate with oxygen set at [−25,0] mmol·gCDW⁻¹·h⁻¹ (based on carbon consumption rate per growth rate) with a fitted linear trendline, GAM is the slope value while NGAM is the intercept value. The fitted slopes for A) all growth rates B) carbon-limited growth rates C) non-carbon limited growth rates are compared. ‘Error bars’ do not show standard deviation; they represent the range of allowed ATP flux for each point.

### S3 Model vs experimental excretion

**Supplementary Figure S2.**
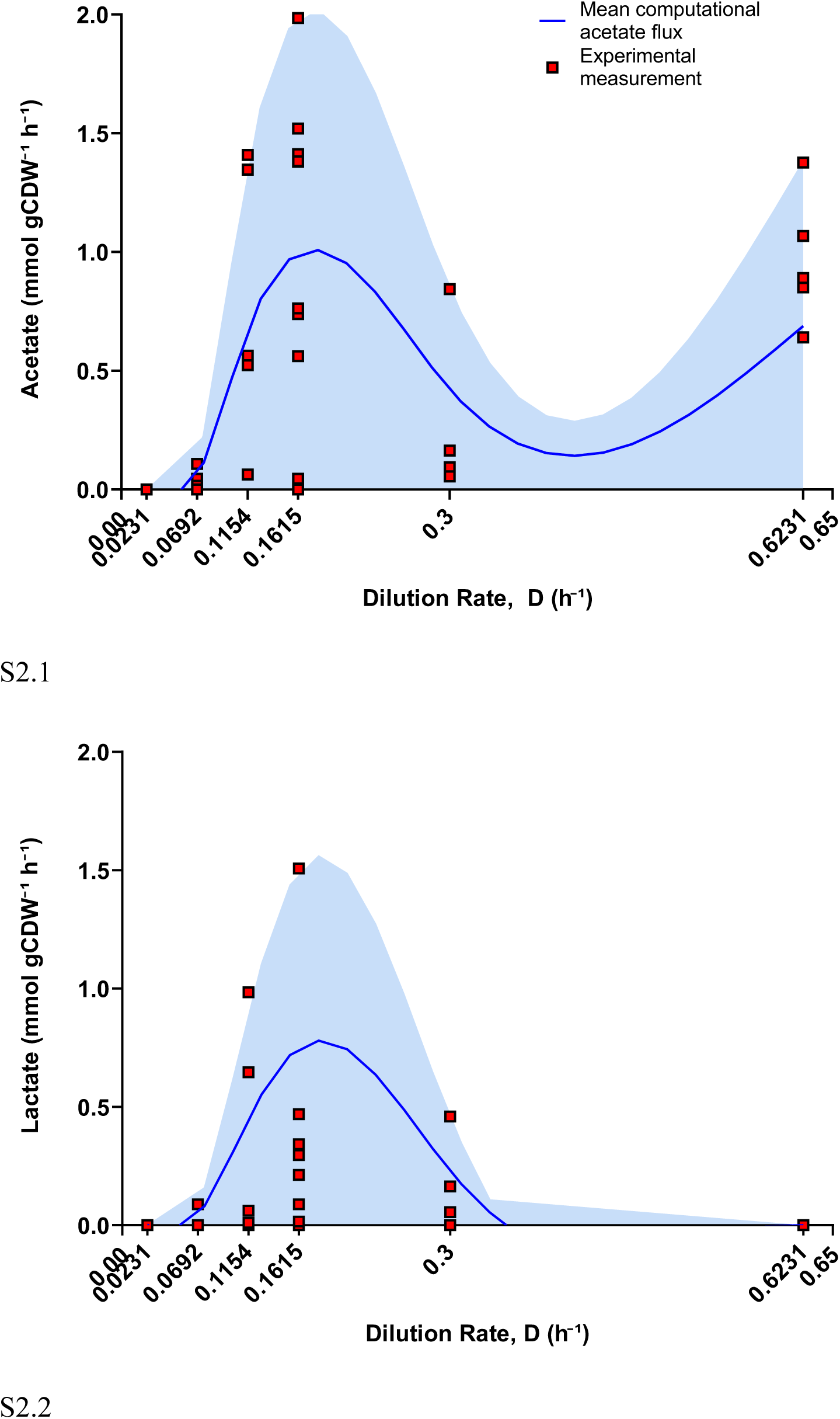
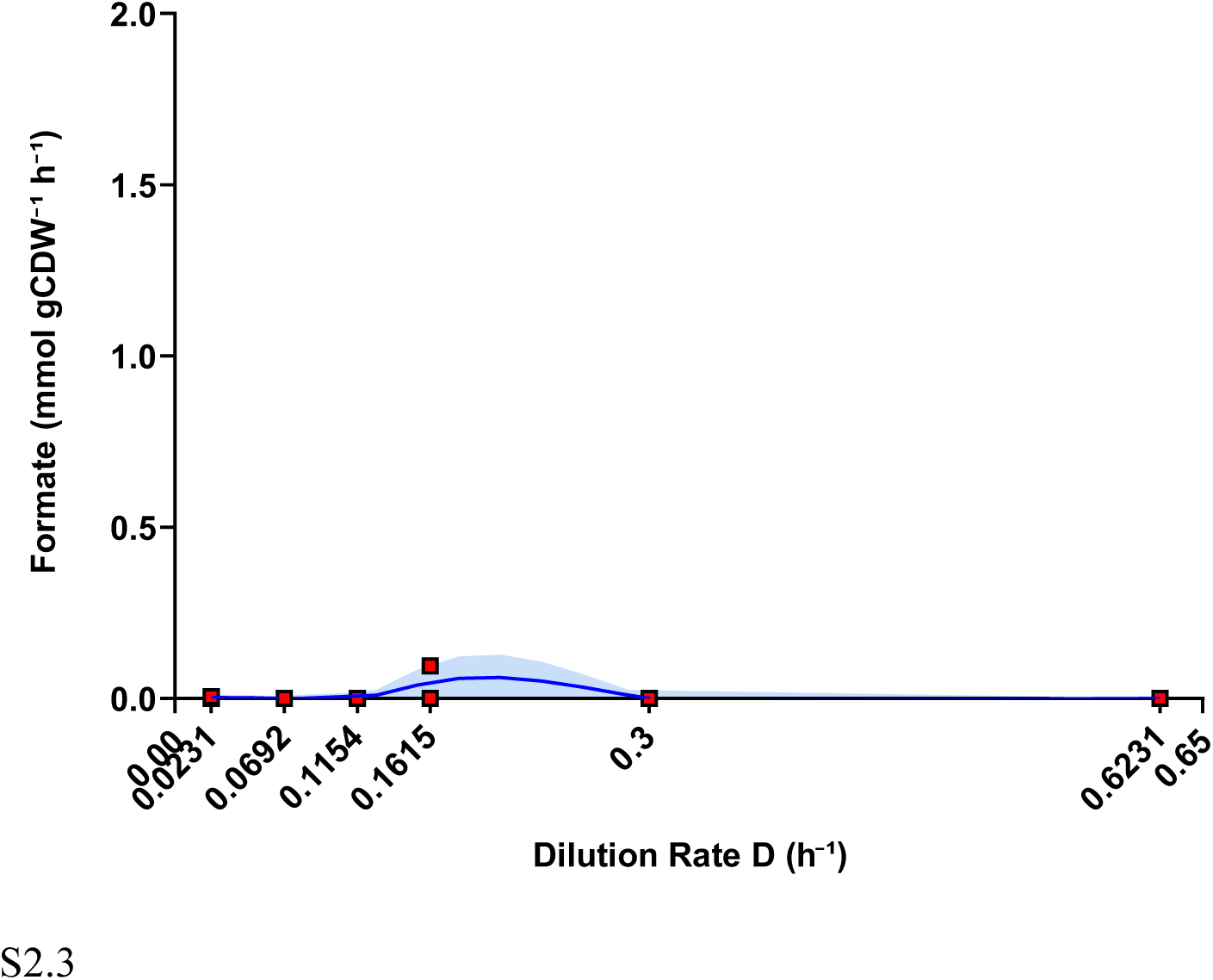
Exploratory comparison of sampled extracellular overflow fluxes with measured chemostat overflow rates. Sampled model distributions for acetate, lactate, and formate exchange were compared with experimental measurements across dilution rates. These panels were used as exploratory checks of extracellular output behaviour and were not used as primary validation evidence because exchange-space constraints and relaxation settings may influence apparent agreement with measured secretion rates.

